# Sclerotic prostate cancer bone metastasis: woven bone lesions with a twist

**DOI:** 10.1101/2023.09.11.557266

**Authors:** Felipe Eltit-Guersetti, Qiong Wang, Naomi Jung, Sheryl Munshan, Dennis Xie, Samuel Xu, Bita Mojtahedzadeh, Danmei Liu, Eva Corey, Lawrence D True, Colm Morrissey, Rizhi Wang, Michael E. Cox

## Abstract

Bone metastasis (BM) are the most severe and prevalent consequences of prostate cancer (PC) affecting more than 80% of patients with advanced PC. PCBM generate pain, pathological fractures, and paralysis. As modern therapies increase survival, more patients are suffering from these catastrophic consequences of PCBM. Radiographically, PCBM are predominantly osteosclerotic, but the mechanisms of abnormal bone formation, and how this “more and new” bone is related to fractures is unclear. In this study, we conducted a comprehensive analysis on a cohort of 76 cadaveric PCBM samples and 12 from non-PC donors as control. We used μ-CT to determine three-dimensional organization and quantify bone characteristics, quantitative backscattering electron microscopy to characterize mineral content and details in bone structure, nano indentation to determine mechanical properties, and we finalize with histological and immunohistochemical analysis of bone structure and composition. We define 4 phenotypes of PCBM, osteolytic, mixed lytic-sclerotic, and two subgroups of osteosclerotic lesions, those with residual trabeculae, and others without residual trabeculae. The osteosclerotic lesions are characterized by the presence of abnormal bone within the trabeculae surfaces and intertrabecular spaces. This abnormal bone is characterized by higher lacunae density, abnormal lacunae morphology and orientation. However, we observed no significant difference between this irregular bone and residual trabeculae in terms of mineral content, hardness, and elastic modulus at micron-scale. The collagen matrix of this abnormal bone presents with irregular organization and is accompanied by increased proteoglycan and phosphorylated glycoprotein content. These characteristics suggests the presence of woven bone in PCBM. However, the lack of subsequent bone remodelling, absence of lamellar bone deposition on its surface, absence of markers of matrix vesicles but evidence of alkaline phosphatase dominated mineralization and collagen-III structure, set up differences from woven bone, while the role of PC cells in inducing this irregular bone phenotype remains unclear.

## 1. INTRODUCTION

Prostate cancer (PC) is the most frequently diagnosed cancer among men, with estimates of ∼1.5 million new cases every year (^1^). Although modern therapies have improved PC survival, about 20% of all PC patients progress to a final incurable metastatic state. Bone metastasis (BM) develops in > 80% of these advanced PC patients (^2^), most frequently involving the trabecular bone of the ribs, epiphyses of long bones, and the vertebral bodies (^3, 4^). PCBM severely affects patient’s quality of life by causing intractable pain, bone weakness and increased risk of fracture. Emerging therapies are extending survival resulting in patients living long enough for BMs to become increasingly relevant (^5^). It is therefore imperative that we better understand the biological mechanisms that cause BMs, and structural changes that they induce. This knowledge is vital for the rational development of effective treatments and palliative care options.

Because of their predominantly osteoblastic nature, PCBM were histologically described as hyperosteoidosis (^6^) or osteomalacia (^7^). Later analysis demonstrated that PCBM exhibits a spectrum of pathologic changes, from pronounced osteopenic to highly osteosclerotic phenotypes (^8^). Because of the lack of collagen matrix alignment in the osteosclerotic regions, the bone of PCBM has been described as “woven bone”. This term is used to describe poorly organized bone, characteristic of fetal bone, and in bone callus formed during the repair process in adults. In both cases woven bone is later replaced by lamellar bone. Although informative, these are relatively low-resolution histologic descriptors, and only a handful of studies have ventured into high-resolution, or three-dimensional analysis of PCBMs. For instance, a series of synchrotron-based high-resolution micro-computer tomography (μCT) scans from two patient samples (^9, 10^), demonstrated alterations in bone mineral density and trabecular morphology and structure, and recently, a high-resolution μCT scan reported similar structural changes in PCBMs obtained from two patients (^11^). These highly relevant descriptions are limited by sample size, and they require further validation to draw definitive conclusions.

Because of the osteosclerotic nature of PCBM, these lesions are thought to be mechanically stable, however, the clinical reality is that PCBM increases the risk of fracture and spinal cord compression, causing severe intractable pain or even paralysis. Despite these critical effects in a large population, very few studies have studied the mechanical properties of bone affected by PCBM. Recently, one study described higher mechanical properties in osteosclerotic metastatic cancer lesions (2 prostate, 2 breast), compared with osteolytic metastatic lesions (1 prostate, 1 breast, 1 lung, 1 kidney), while another study described higher vertebral strength in osteosclerotic metastatic lesions (1 prostate, 7 breast, 2 esophageal, 3 lung, 8 kidney), than mixed (1 prostate, 4 breast, 1 esophageal, 3 lung) or osteolytic (1 prostate, 1 breast, 1 esophageal, 3 lung). Neither of these studies tested healthy control samples, to establish conclusions on the fracture risk of sclerotic lesions (^12^). To the best of our knowledge, a total of only about a dozen PCBM lesions have ever been structurally analyzed at high (micron) resolution or for their mechanical properties. Clinically, increased fracture risk in PC patients is linked to lower bone density due to hormone ablation therapy (^13^), while in male multiple myeloma patients, vertebral fractures are associated with trabecular thickening and sclerosis of three or more vertebrae, suggesting an association between fractures and osteosclerosis (^14^).

Here, we aimed to perform a comprehensive investigation into the spectrum of alterations in composition, structure, and mechanical properties of vertebrae affected by PCBM, hypothesizing that alterations in the structural organization of bone induced by PC increases fracture risk. To achieve this goal, we use multiple high-resolution and three-dimensional techniques to analyze a cohort of 76 cadaveric PCBM samples from 43 men and 12 samples from healthy donors as control. The microstructure was analyzed by mCT. High resolution morphology, Ca wt%, properties of cellular lacunae was investigated through backscattered imaging in scanning electron microscopy. We explored the alignment of collagen in bone and complemented the analysis of the extracellular matrix composition by immunohistochemistry. We observed a high variation in structure and mineral content among PCBMs, that could be grouped in distinct phenotypes. Osteosclerotic PCBM are characterized by the presence of deposition of irregular mineralized matrix in the intertrabecular spaces, which present increased lacunae density and size as well as poor collagen organization. This matrix also shows an increased content of collagen 3, mineralization related proteins and proteoglycans, suggesting differences in the osteogenic process. Our findings offer valuable insights and foundational information into the characteristics of PCBM, while they have the potential to serve as a basis for investigating the mechanisms behind their formation and their clinical implications.

## 2. MATERIALS AND METHODS

### 2.1 Cadaveric vertebral metastasis of PC samples

All the described procedures were approved by the institutional review board at the University of Washington (2341) and the University of British Columbia (H21-02668). We obtained cadaveric samples (n=76) from the autopsies of 43 patients who died of metastatic CRPC and signed written informed consent for a rapid autopsy to be performed ideally within 2 hours of death, under the Prostate Cancer Donor Program at the University of Washington Medical Center (^15^). As control samples, we obtained 12 vertebrae from 3 age matched male donors from the University of British Columbia Body Donation Program. The demographic and clinical information of the donors is provided in Table <u>S1</u>. The specimens were obtained by using a 11 mm diameter trephine and stored in 70% ethanol until analyzed with a μCT scanner.

### 2.2 Three-dimensional μCT scanning of cadaveric PC bone specimens

To analyze the mineral structure of cadaveric vertebral samples, we used a μCT scanner (Scanco Medical, Bruttisellen, Switzerland) with the following setting: 70 kVp, 114 mA, 8 W, 500 ms integration time. A total of 1500 slices with an isotropic voxel size of 10μm were obtained on each specimen according to the manufacturer’s standard protocol. We extracted the following bone parameters; trabecular bone volume/total volume (BV/ TV), trabecular number (Tb.N, mm1), trabecular thickness (Tb.Th, mm), trabecular separation (Tb.Sp, mm), Connectivity Density (Conn-Dens), and additionally, we used an automated segmentation algorithm (Image Processing Language, Version 5.08b, Scanco Medical) to determine the bone mineral density (BMD, mg HA/cm3). To avoid considering the damage and debris created by obtaining the samples, a perimetric volume 0.7mm from the edges was removed from each sample prior analysis. Areas with evidence of cortical bone were also excluded.

For qualitative analysis, we analyzed three 30-micron thick CT-slides (3 voxels in depth) on each sample, one in the boundary between the top ¼ and the following ¼, one in the middle of the sample, and one in the boundary between the 3rd ¼ and the bottom ¼. We then assessed tomograms for the presence and distribution of trabeculae to define PCBM phenotypes defined in results (Section 3.2).

### 2.3 Quantitative Backscattered Electron-Scanning Electron Microscopy analysis on cadaveric PC bone specimens

After mCT scanning, the cadaveric specimens were transversally cut with a water-cooled diamond saw (IsoMet 4000, Buehler, Lake Bluff, IL, USA), into two sections corresponding to 1/3 and 2/3 of the total length of the sample. The smaller bone section of each sample was used for histology (section 2.6), while the larger ones were immersed in a sequence of acetone (70%, 90% and 100% x2) for dehydration (2 days in each concentration). The dehydrated bone sections were then infiltrated in 50%, 80% and 100% x2 embedding medium (Spurr low viscosity kit, PELCO, USA) in acetone for 3 days in each concentration. After infiltration, bone sections were embedded together with Carbon (Spec-Pure Carbon Rod, TED PELLA) and Aluminum (99.99% Al, TED PELLA) for quantitative backscattered electron analysis. The transverse surface of the embedded bone sections underwent grinding using a series of decreasing grit carbide papers, and subsequently polished using a 1 µm diamond suspension. Following this preparation, the surface was coated with carbon and subjected to scanning electron microscopy (SEM, FEI Quanta 650, Oregon, USA) imaging in backscattered electron (BSE) mode. The BSE images were captured at a working distance of 15 mm with a magnification of 200×, resulting in image dimensions of 636 µm × 457 µm and a resolution of 1536 pixels × 1326 pixels. An equal number of randomly-selected areas of interest representing distinct bone morphologies in each specimen were selected and the mineral density was determined for these designated areas and expressed as Ca wt% (peak value representing the most prevalent calcium concentration) following the methodology outlined by Roschger et al. (1998) (^16^).

### 2.4 Nanoindentation

We employed a nanoindentation system (Nano Indenter XP System, MTS Nano Instruments, Oak Ridge, TN, USA) featuring a Berkovich tip to assess the elastic modulus and hardness of the embedded bone samples. Prior to the measurements, the area function in the nanoindentation system underwent calibration using the Young’s modulus of fused silica (72 GPa). Additionally, calibration of the indenter tip relative to the microscope was performed before initiating the indentation process. The measurements were conducted on the polished embedded bone sections, ensuring a controlled penetration depth of 1.0 µm. The loading strain rate was set at 0.05 s-1, with a holding time at the peak load of 10 seconds. Subsequently, the tip was unloaded to 90% of its peak load. The elastic modulus and hardness values were derived from the load versus displacement curve utilizing the Oliver and Pharr method (^17^).

### 2.5 Lacunae analysis

Based on the analysis of BSE images, we selected 10 areas of interest within each of the 12 samples, ensuring representation across various lesion types, including osteosclerotic, mixed, and osteolytic. Within each image (636 µm × 457 µm, 1536 pixels × 1326 pixels), we determined areas of lamellar bone and disorganized bone (sclerotic lesions) based on the presence or absence of a lamellar structure as previously described. We then extracted essential information concerning lacunae density, lacunae dimensions, lacunae area relative to bone area, and lacunae orientation. The results of lacunae properties were obtained through the MATLAB script through multiple iterations in the following steps: The quantitative BSE (qBSE)-SEM image is imported into the program, converted to grayscale, cropped to remove the info bar, and a gaussian filter is applied. The grayscale threshold (corresponding to 8 wt% Ca) is determined to create a mask of the pores present in the bone. Pores with an area between 2.5 µm^2^ and 500 µm^2^ are identified as lacunae and designated with an ID. The area, centroid, major axis length, minor axis length, ratio between axes, circularity, eccentricity, orientation (compared to horizontal) is derived from each lacuna. We manually excluded cracks and artifacts. We then quantified the average lacunae size, total lacunae area, average lacunae size, total region of interest rea, number of lacunae, lacunae density, and the lacunae area/bone area fraction for every image.

### 2.6 Histology and immunohistochemistry

Sections obtained from the cadaveric specimens were fixed in 10% buffered formalin for 24 hr, rinsed in phosphate-buffered saline (PBS) three times for 1 hr each, and decalcified in 10% formic acid for 5 days. The specimens were then dehydrated in an ethanol series (70%, 80%, 90%, 95%, 100%), cleared in xylene substitute, and paraffin embedded. Serial 5 µm thickness sections were prepared and mounted on (name and course of product) microscope slides. One slide of each core serial section was stained with hematoxylin-eosin, picrosirius red, toluidine blue and Goldners’ trichrome, respectively.

We used Picrosirius red staining to quantify birefringence as an indirect method of measuring collagen alignment (^18^). Images including ∼30% of the histological sections were taken under bright field and polarized light. Pictures with a field of view of 6 x 6 mm were obtained using low magnification in the centre of the tissue section. We used ImageJ software to quantify the bone area, by measuring the percentage covered by bone in each image using the red channel in pictures taken bright field. The birefringence was calculated by measuring the area detected by the green channel polarized light images.

For immunohistochemical analysis, we treated mounted sections with 2% pepsin in 0.1N HCl for 20 min at 37 degrees for antigen retrieval, followed by endogenous peroxide quenching using 3% H_2_O_2_ for 15 minutes. For non-specific protein blocking we used 3% BSA in PBS for 20 min. We incubated the samples with primary antibodies (list of antibodies in Table S2) overnight at 4 °C in a humidified chamber. After washing with PBS, we incubated the samples in HRP-conjugated secondary antibodies (Table S2) for one hour at room temperature. We revealed the immune detection using a commercial detection kit (VECTOR) following manufacturer instructions. Observations were performed using an upright microscope (Zeiss Axiophot). Images were obtained with a cooled camera and processed and collected using ZEN software (Carl Zeiss Canada; North York, ON). Post processing of images was performed using Adobe Photoshop.

## 3. RESULTS

### 3.1 PCBM are variable, predominantly osteosclerotic and some osteolytic, characterized by increased or decreased trabecular number respectively

Previous histological assessments from the same biobank characterized vertebral body PCBM lesions as exhibiting a spectrum of phenotypes ranging from predominantly osteoblastic to mostly osteolytic (^19^). These results were based on a limited number of thin sections of a specific region of each vertebral core specimen. To evaluate the overall characteristics of the lesion, we performed quantitative μCT analysis of the whole core obtained from prostate cancer-involved vertebral samples (Tables S3). We observed that of the 76 analyzed samples, 55 exhibited increased BV/TV compared to our age matched control group (mean ± St. Dev.). In contrast, only 9 samples exhibited decreased BV/TV relative to the control group (Fig 1A).

**Figure 1:**
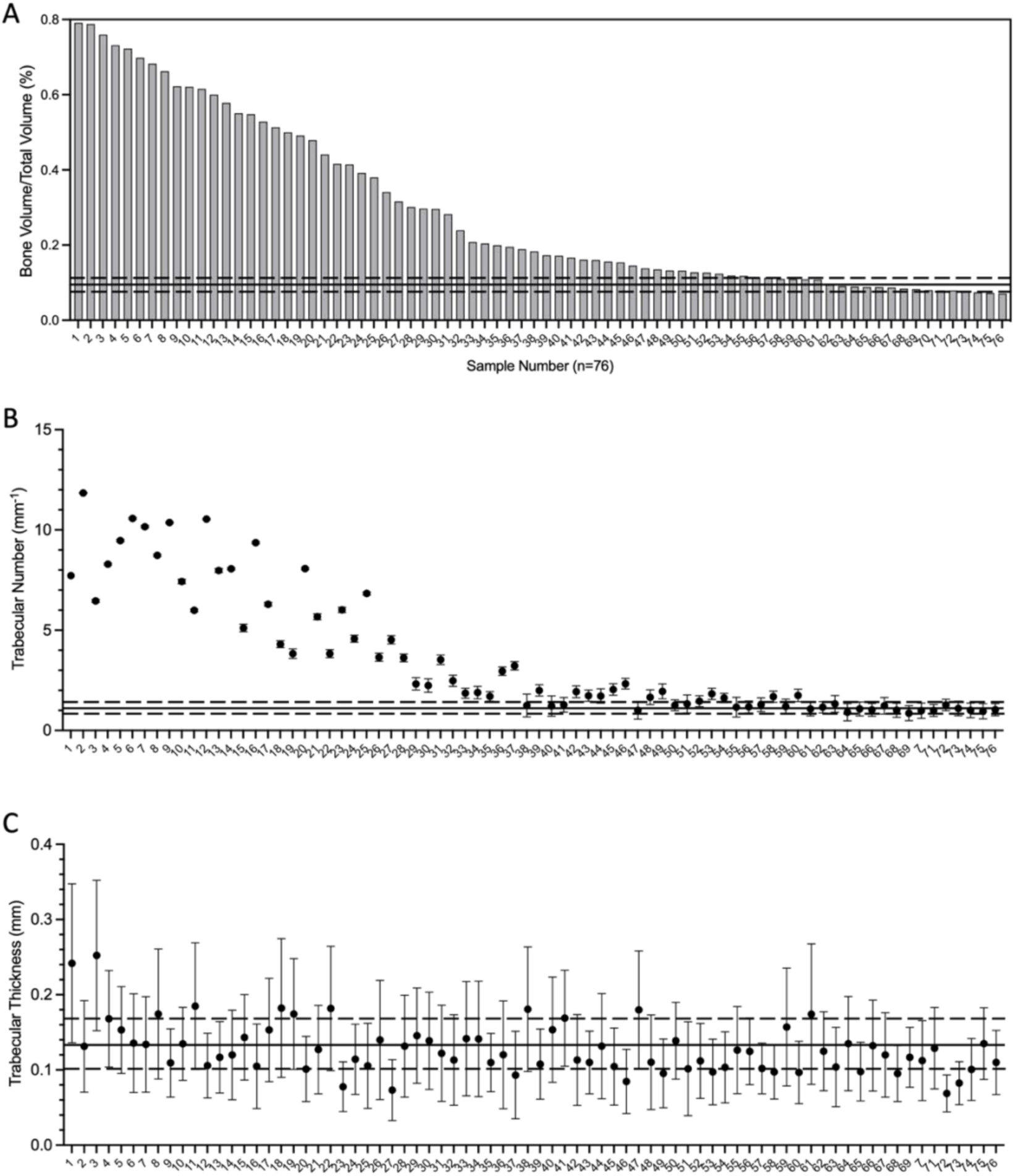
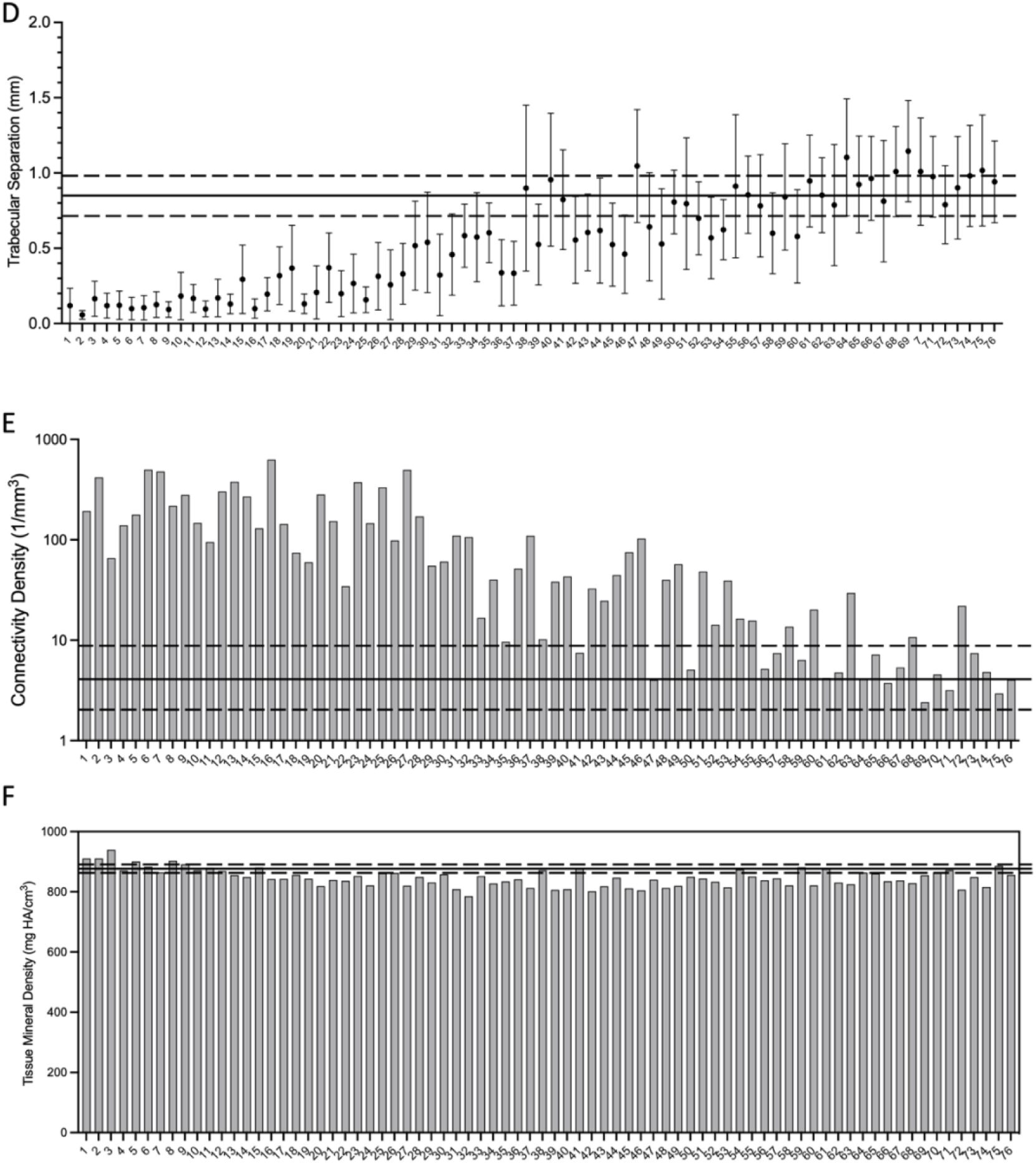
Micro-CT analysis of cadaveric vertebral prostate cancer bone metastasis (n=76). A) Waterfall plot ranking by bone volume over total volume of PCBM samples. B) Trabecular number by sq mm, mean value and SEM of each sample is shown. C) Trabecular thickness in mm, mean +-standard deviation is shown. D) intertrabecular space in mm, mean +-standard deviation is shown. E) Mean trabeculae connectivity density (number of connections per trabeculae) in PCBM samples (Chart in Log scale). F) Mineral density of the bone fraction (excluding medullary spaces) of PCBM samples, expressed in mg of hydroxyapatite/ cubic mm. Solid line, mean value and dashed lines +-St. dev. of 12 age matched vertebral control samples.

Additionally, 55 of the samples have increased Tb.N compared to controls, and only 3 have decreased Tb.N (Fig 1B). However, this increase in Tb.N is not related to Tb.Th which, on average, were indistinguishable from the controls, but that exhibited a larger variation within and between samples (Fig. 1C). As expected, increased Tb.N is accompanied by decreased Tb.Sp, and increased Conn-Dens (Fig. 1 D-E). We next assessed radiodensity of the mineral bone fraction of the specimens and observed that PCBM exhibited lower BMD than the control group (Fig 1F). These results validate the consensus that most PCBM are predominantly osteoblastic lesions, with increased volume of mineralized tissue, mostly due to an increase in the number of trabeculae, but no change in their thickness. A smaller group of lesions (∼15%) are predominantly osteolytic with decreased BV/TV, characterized by fewer trabeculae, rather than thinner trabeculae.

### 3.2 PCBM have complex structures exhibiting areas of osteolysis, as well as osteosclerosis dominated by a disorganized matrix in the presence or absence of residual lamellar trabeculae

μCT analysis allows for a complete 3-dimensional reconstruction of the vertebral core sample. Combining a conventional spectrum of μCT measurements of trabecular bone with a qualitative analysis, we observed differences in mineralized matrix architecture in PCBM specimens that are generally consistent with those previously described in this cohort of specimens by histologic methods (^8^). Based on qualitative observation, we define four distinct dysmorphic bone patterns in PCBM (Fig. 2) which are described as follows. Osteolytic samples (n=16) are those with at least 3 voids larger than 1 mm without trabeculae, and the presence of thinned or broken trabecula (Fig 2B). Mixed samples (n=20) exhibit both osteolytic and osteosclerotic areas (Fig 2C). Osteosclerotic with residual trabeculae (n = 23) are those in which we observed more than 5 trabeculae (each larger than 1 mm diameter) in each section accompanied by increased trabecular thickness and the presence of intertrabecular mineral material (Fig 2D). Osteosclerotic without residual trabeculae (n = 15) have less than 5 trabeculae per section and dominated by accumulation of irregular radio-dense material (Fig. 2E).

**Figure 2:**
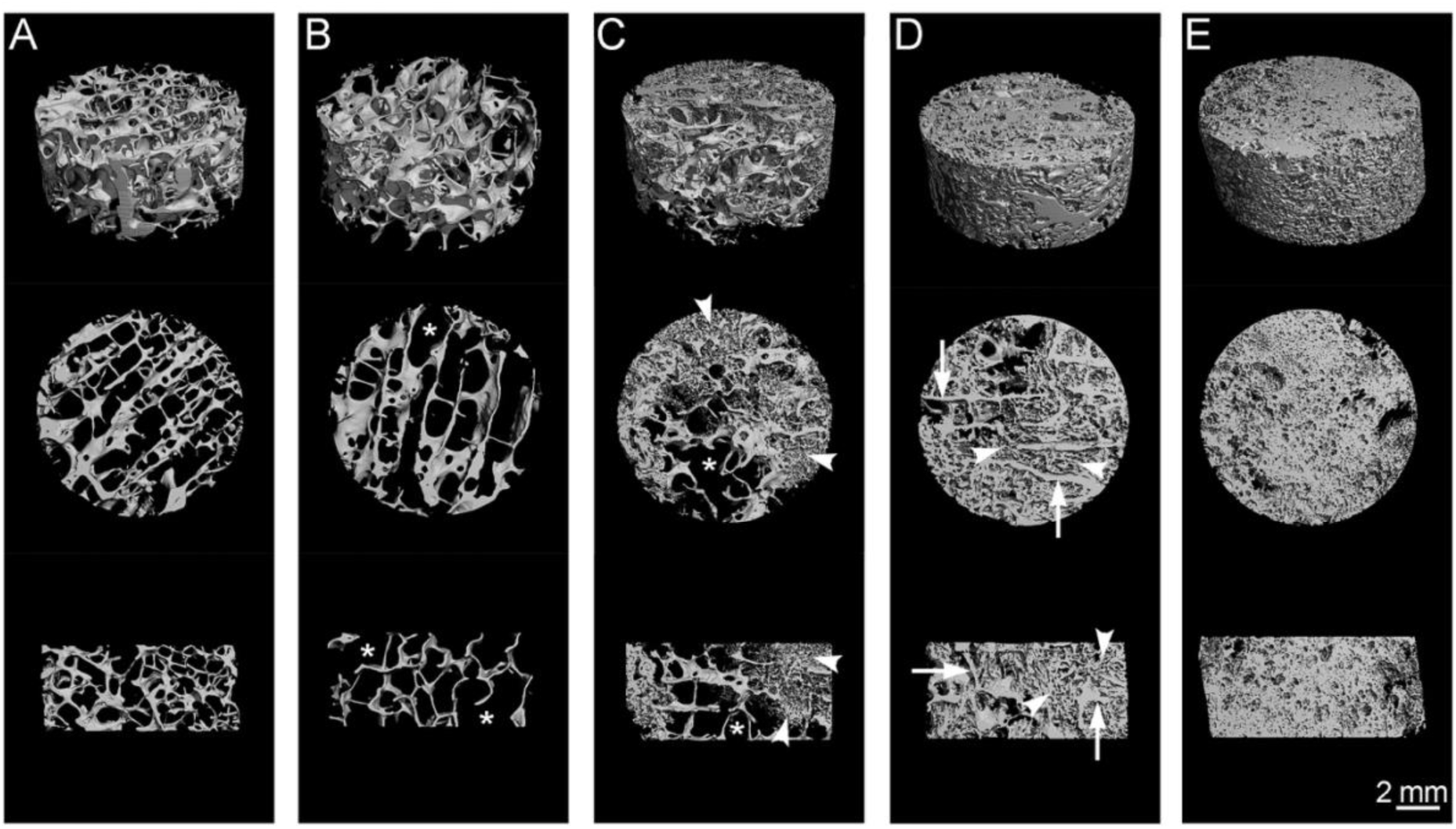
Renders of representative cadaveric vertebrae of every sub-type obtained from Micro-CT images. A) cadaveric age-matched control, B) osteolytic sample with large voids (Asterisks) and thin-broken trabeculae (arrowhead), C) Osteoblastic PCBM sample, with residual trabeculae (arrows), which are thicker, and characterized by deposition of mineral in the intertrabecular spaces (arrowheads). E) Osteoblastic sample without residual trabeculae, showing a homogeneous matrix of irregular mineral deposition along the whole sample. Top views = 4.5 mm thick sections, middle and bottom sections = transverse view and sagittal view of 1 mm thick sections. Renders were generated by superimposition of 6-micron sections.

We then compared the quantitative values provided by μCT with the qualitative classification of our morphological descriptions, we found that the μCT values are concordant with the differential structural pattern (Fig 3A, Table S4). When comparing specific parameters, we observed that the osteosclerotic samples with few or no detectable residual trabeculae (Os WoT) had an average BV/TV ratio that was significantly higher than that of the osteosclerotic group with residual trabeculae (Os Tra), which not surprisingly is higher than Mixed, Osteolytic and control samples (Fig 3A). As expected, we observed the lowest Tb.Sp in the osteosclerotic groups compared with the mixed samples, and the highest values in the osteolytic samples (Fig 3B). The Os WoT also exhibited a higher Tb.N than the Os Tra, which in turn is higher than that of the Mixed and Osteolytic (Fig 3C). While the Conn-Dens was indistinguishable between the osteosclerotic groups, their Conn-Dens was greater than that of the Mixed, and these exhibited a greater Conn-Dens value than osteolytic and control samples (Fig 3D). Surprisingly, we observed a higher BMD density in the Ob WoT and control specimens compared to other groups, which in turn, were indistinguishable (fig 3E), and that Tb.Th was indistinguishable between all the groups (Fig. 3F). Combined, these results distinguish osteoblastic lesion phenotypes depending on the presence, or lack, of residual trabeculae, that are independently evident by quantitative μCT. These results also suggest a higher mineral density in the Os WoT and the control group, suggestive of different matrix composition.

**Figure 3:**
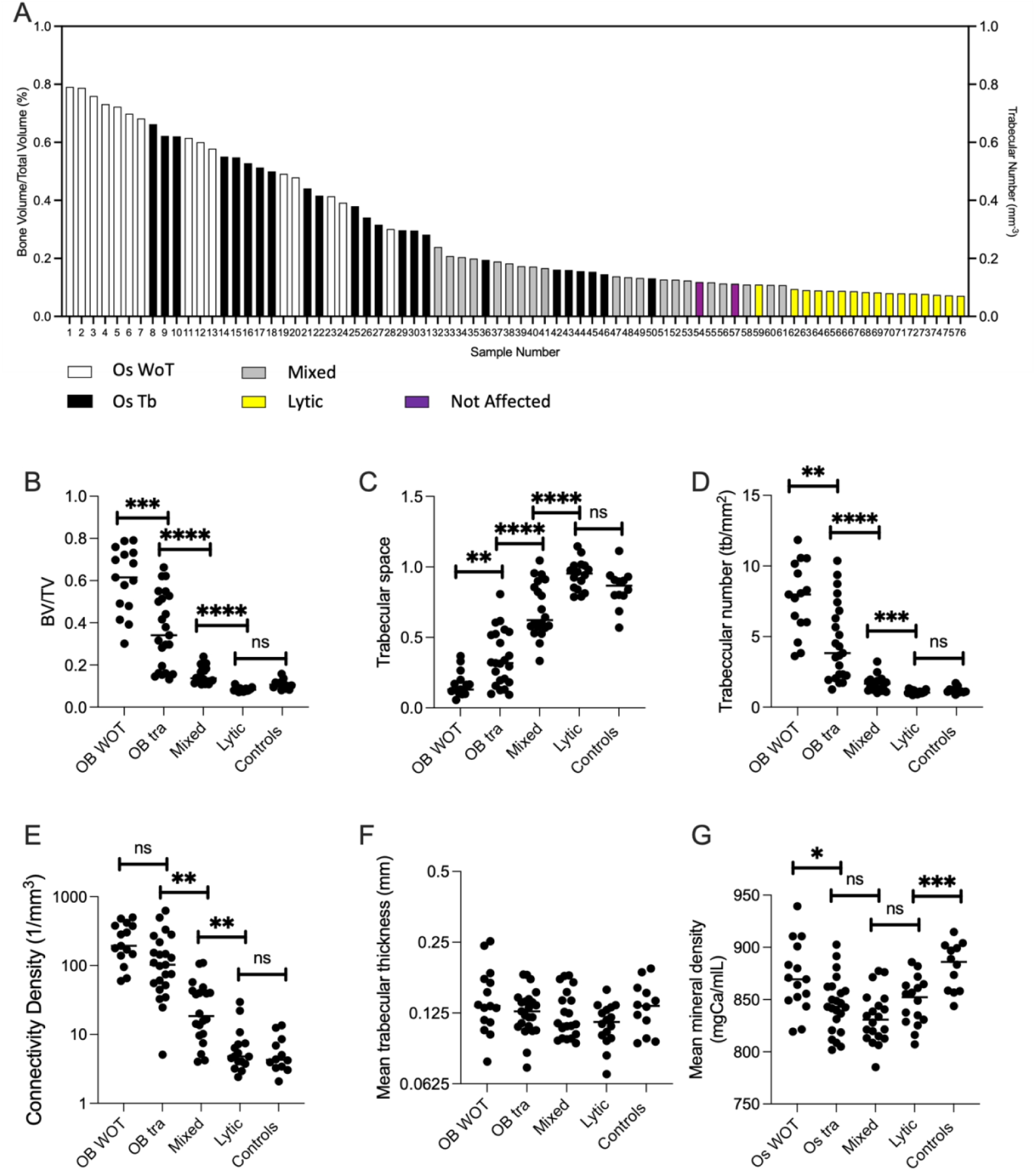
Quantitative analysis of samples based on lesions phenotype. A) waterfall plot of bone volume / total volume, of samples based on phenotypical presentation of micro-CT measurement. B-G) Dot plots of quantification analysis of different sample phenotypes according to micro-CT evaluation.

### 3.3 Different PCBM phenotypes are present within a patient

To evaluate whether PCBM within a patient present with similar morphologic features, we analyzed samples from the 21 patients from whom we have 2 or 3 vertebral core samples using the previously described parameters. We first compared our qualitative phenotypic description for those samples and observed that 16 of these patients presented with variable phenotypes, while only 5 patients had two or more samples with similar phenotype (Table S5). To quantitative evaluate the similarities, we compared the range of the BV/TV within a patient with the mean value of BV/TV for that patient. We observed that 8 of the 21 samples had a BV/TV range that was less than 20% of the patient BV/TV average, while 13 have BV/TV values that were greater than 20% of the patient’s average BV/TV ratio (Fig. 4). Since our samples represent just a limited number of PCBM existing in a patient, the presence of diversity in phenotypes and BV/TV suggest that patients present different types of PCBM.

**Figure 4:**
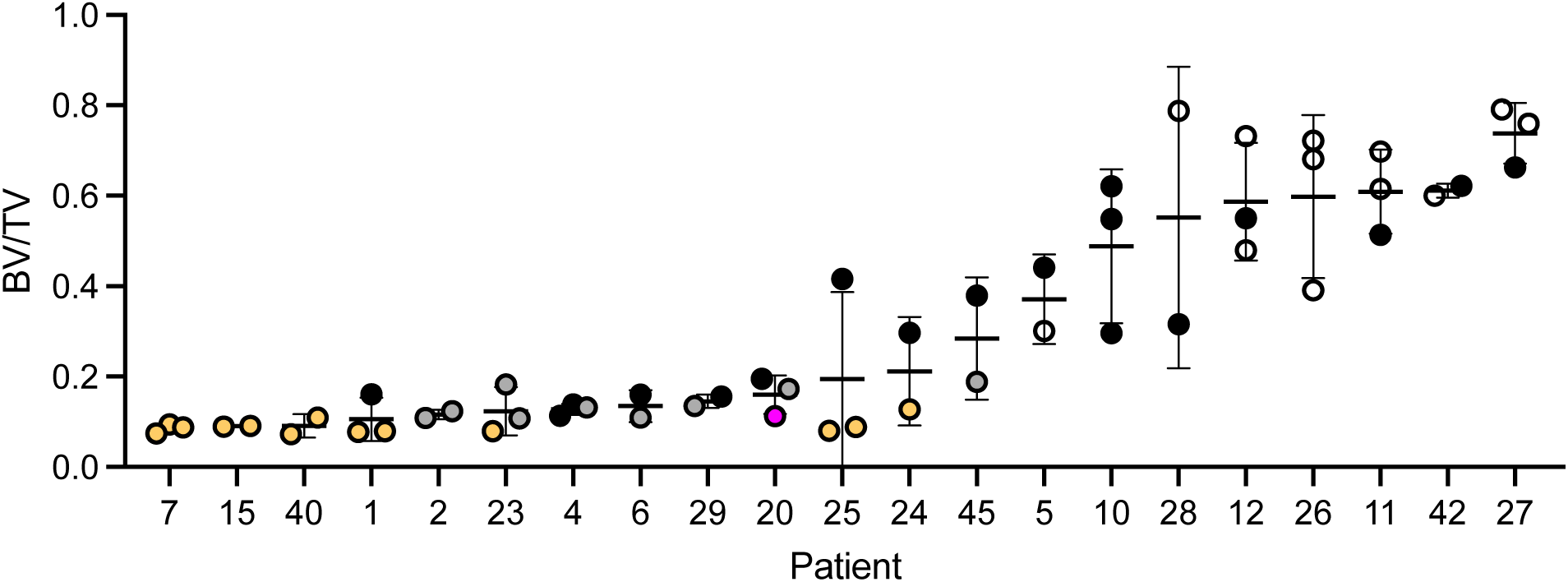
BV/TV of samples grouped by patient. The patients number corresponds with the numeratin assigned in table S1. Color code: Yellow = Lytic, grey Mixed, Black= Os Tra, white= Os WoT.

### 3.4 Irregular bone deposits on the surface of pre-existing trabeculae and occupies the intertrabecular spaces in osteosclerotic PCBM

To gain deeper insights into the microstructure of the PCBM vertebral specimens, we conducted high-resolution imaging analysis using a scanning electron microscope (SEM). We focused on 12 PCBM which were samples selected to represent the spectrum of PCBM presentations based on BV/TV and the qualitative observations previously described (Fig. S1). Representative examples of the observed morphological features are shown in Figure 5.

**Figure 5:**
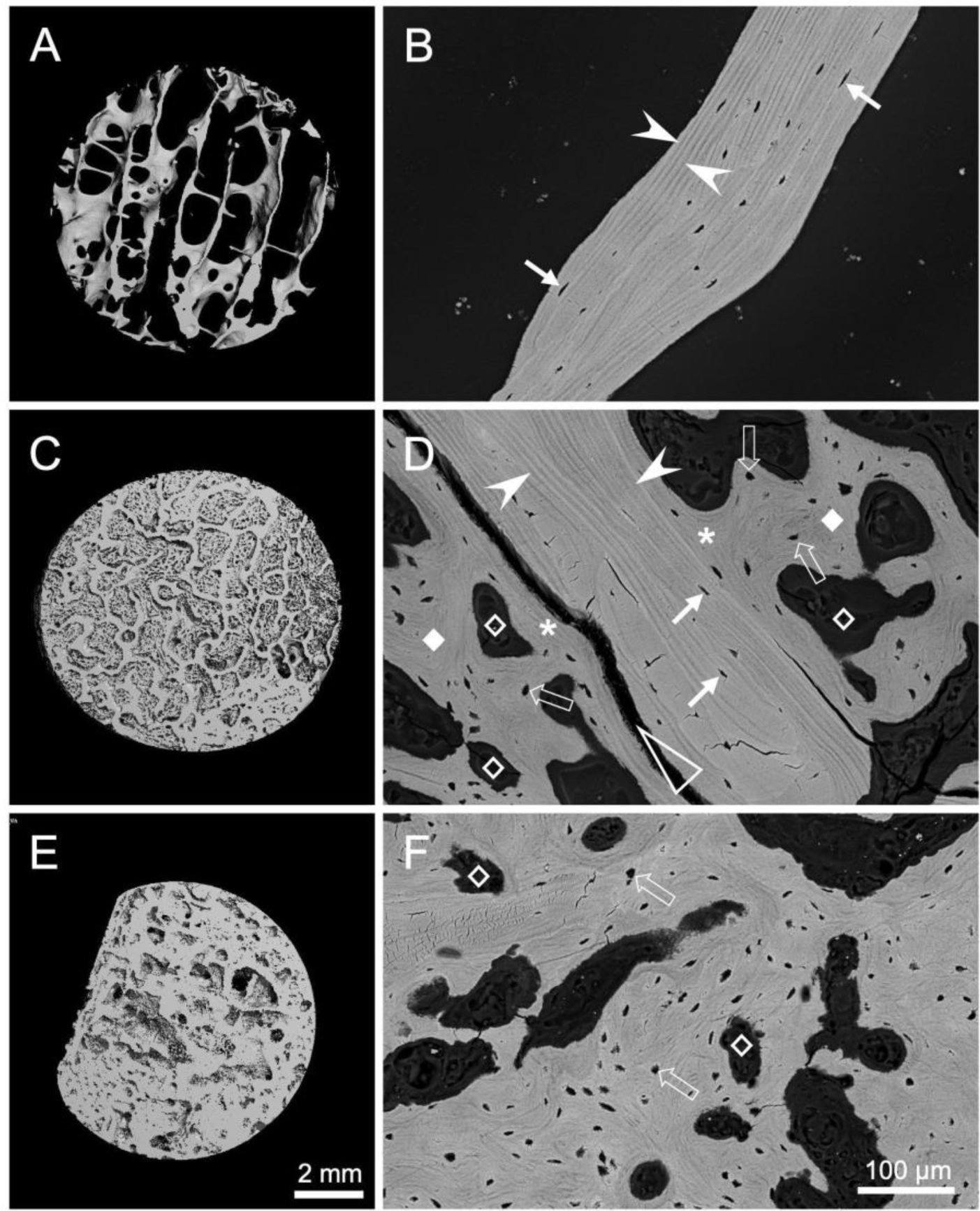
Backscattered electron microscopy observation of features observed in PCBM. A) micro-CT image of an osteolytic PCBM. B) Backscattered SEM observation of sample in A. lamellar structure is observed as light and dark bands (arrowheads) that align parallel to the major axis of the trabeculae. Lacunae are observed as lentil-shaped voids (arrows) aligned parallel to the lamellae orientation. C) micro-CT image of sample of osteoblastic with pre-existing trabeculae PCBM. D) Backscattered SEM image of the sample in C. pre-existing trabeculae is evident by the presence of lamellar structure with light and dark bands (arrowheads), and the presence of lacunae with their major axis oriented parallel to the orientation of the lamellae (arrows). Osteosclerotic irregular bone is evident on the surface of the pre-existing trabeculae (asterisks), and it extends towards the intertrabecular space (diamonds). This irregular bone has no evidence of lamellar structure, the lacunae do not follow any specific orientation, and many of them are even round-shaped (open arrows). The presence of large voids is noticeable (open diamonds). The interface between the pre-existing trabeculae and the osteosclerotic bone is usually visualized by the presence of a crack between the mineralized tissues (open arrowhead). E) micro-CT image of sample of osteoblastic without pre-existing trabeculae PCBM. F) Backscattered SEM image of the sample in E. the space is completely occupy by irregular osteosclerotic bone, with absence of lamellar structure, lacunae with no defined orientation (open arrows) and large voids (open diamonds). A, C, E) images generated by superposition of images with a voxel size of 6 um to obtain a 1 mm thickness image.

In osteolytic PCBM samples (Fig. 5A-B), the trabeculae displayed a characteristic lamellar structure, denoted by striations along the major axis of the trabecula, indicative of lamellar organization (Fig 5B arrowheads). Lacunae in these regions were elongated, with their major axis parallel to the lamellae (Fig 5B arrows). in Osteosclerotic samples with residual trabeculae (Fig. 5C-D), the presence of persisting trabeculae was discernible by the lamellar matrix (Fig 5D, arrowheads) and the anisotropic lacunae (Fig. 5D arrows). However, an abnormal, non-trabecular mineralized matrix was also evident on the surface of residual trabeculae (Fig. 5D asterisks). This atypical, mineralized matrix is a hallmark of “sclerotic” regions in PCBM (^8^). It is characterized by the absence of lamellar lines, irregularly shaped lacunae (open arrows in Fig. 5D), and the presence of large, irregular voids exceeding the size of normal lacunae (open diamonds in Fig. 5D). This irregular sclerotic bone protruded from the trabecular surface into, and often occupied most of, the intertrabecular/medullary space (diamonds in Fig. 5D). Notably, distinct cracks were often observed between the sclerotic bone and the trabeculae in SEM imaging (open arrowhead in Fig. 5D), possibly representing the boundary between trabeculae and sclerotic bone. In contrast, the osteosclerotic lesions without residual trabeculae lacked regions of anisotropic trabeculae and were entirely composed of irregular matrix (Fig. 5E-F), characterized by irregularly shaped lacunae and voids, similar to those described in figure 5D. (Fig. 5F). These observations suggest a differential composition of the osteosclerotic PCBM bone than trabecular bone, characterized by absence of trabecular organization and irregular cellular organization.

### 3.5 Unaltered mineral content is associated with normal microscale hardness and elastic modulus in osteosclerotic PC bone

Given the distinct morphological characteristics of the sclerotic lesions, which exhibit differential backscattered electron density compared to adjacent trabecular structures (Fig. 4D), coupled with our prior observation of varying mineral density in different PCBM types (Fig 3G), we postulated the presence of differing mineral content between irregular PC-associated sclerotic bone and residual trabeculae. To test this hypothesis, we conducted a qBSE-SEM analysis to assess the calcium content of the 12 specimens imaged with SEM. In each specimen, we measured at least 5 and up to 9 areas of both sclerotic and trabecular bone. Our findings revealed that the measured calcium weight percentage (Ca wt%) ranged from 21 to 25 Ca wt% across all specimens and regions. While some variations in Ca wt% were observed within specimens between sclerotic and trabecular bone, our analysis showed no significant difference in calcium content between the sclerotic and trabecular bone across the examined regions of the 12 specimens (Fig 6A). These results although limited by sample number and area of analysis, suggests that there no dramatic differences in the mineral content between osteosclerotic bone and trabecular bone.

**Figure 6:**
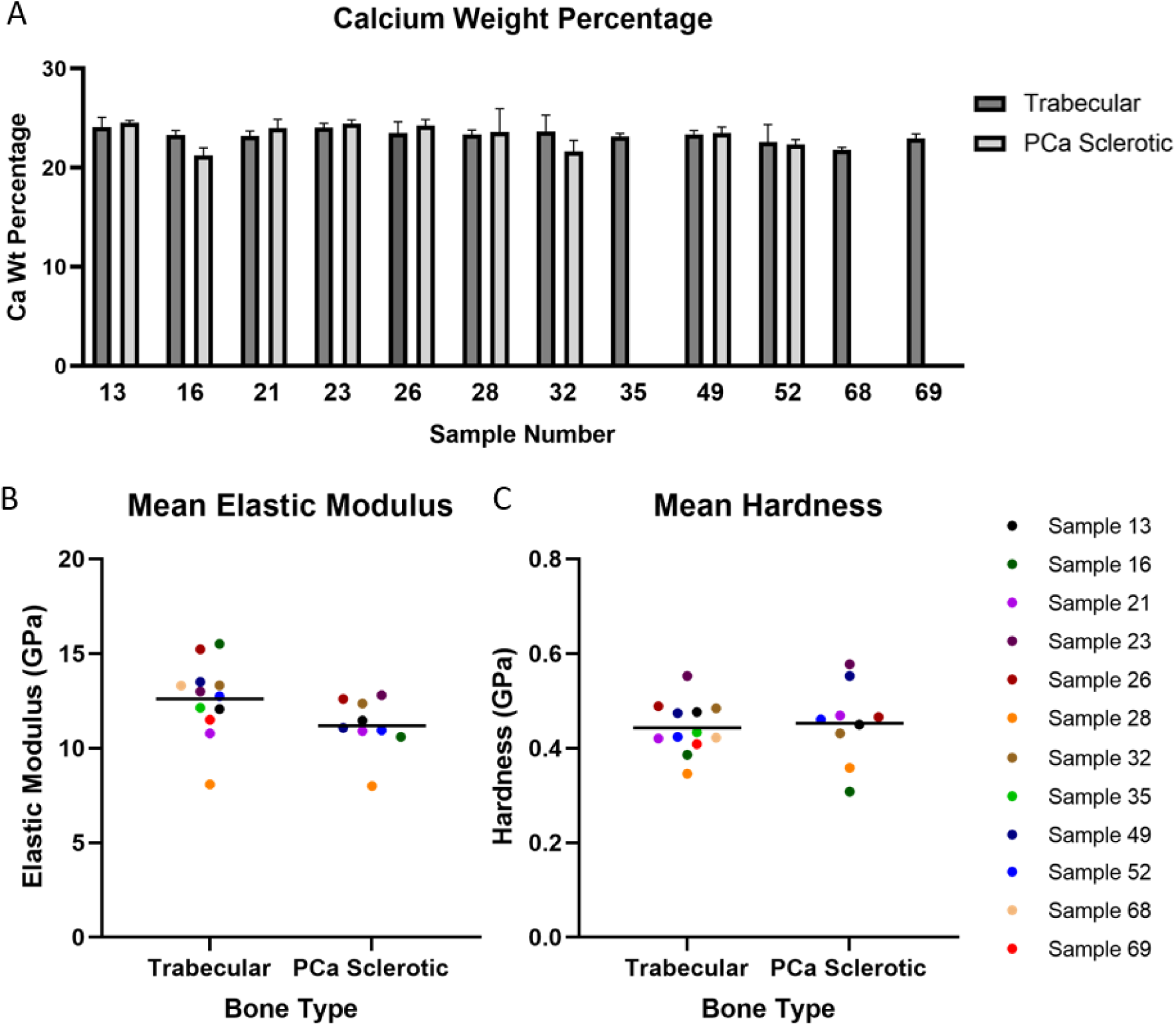
Mineral content and nanomechanical properties of PCBM. A) Calcium weight percentage of cadaveric vertebral prostate cancer bone metastasis samples measured by quantitative backscattered electron SEM. The mean value and standard deviation of the median Calcium weight percentage of each image is shown (images per sample per bone type = 5-9). B) Elastic Modulus and C) Hardness measurements for cadaveric vertebral prostate cancer metastasis samples (n=12). The mean value for each sample in trabecular regions (indents per sample=14-23) and sclerotic regions (indents per sample=7-23) are shown as individual dots. Black lines represent the mean values for all samples.

The mechanical properties (hardness and elastic modulus) of bone are related to the clinical fractures and pain (^20^). To characterize the mechanical properties of the irregular PC-associated bone, we conducted nanoindentation tests at the micron level, following established protocols (^21^) (Fig 6B-C). For each specimen, we calculated the mean from a minimum of 10 measurements within trabecular and sclerotic regions, respectively, for hardness and elastic modulus. Our results indicated that in the trabecular regions, the elastic modulus ranged from 7 to 16 GPa, with a mean value of 12.33 GPa, while in the sclerotic regions, the elastic modulus ranged from 7 to 13 GPa, with a mean value of 11.19 GPa (Fig. 5B). Notably, the elastic modulus between the trabecular and sclerotic regions was indistinguishable (p value = 0.13).

Furthermore, the elastic modulus values of these PCBM specimens closely resemble those of normal trabecular bone, as reported in the literature (^22^). Meanwhile, the measured hardness values between trabecular and sclerotic regions exhibited no discernible distinctions within or across the cancer-involved vertebral bone core specimens. These values also aligned with those of normal trabecular bone. Given that our nanoindentation analysis included all the locations where we measured Ca wt%, we explored potential association between hardness, elastic modulus, and calcium mineral content at each measured site. However, our analysis revealed no association between calcium content and these mechanical properties (Fig S2). In conclusion, despite noticeable morphological changes, our findings suggest that PCBM sclerotic bone possesses mineralization, hardness, and elastic modulus levels that are indistinguishable from those of trabecular bone.

### 3.6 Higher lacunae density, abnormal lacunae morphology and orientation characterize osteosclerotic bone in PCBM

Lacunae house osteocytes which are critical to bone modelling. Lacunae are also the sites mechanically concentrating stress, favoring crack initiation and fracture development (^23, 24^). In our investigation, we examined the lacunar properties within irregular PC-associated bone to ascertain how these lacunae differ from those within trabecular bone. Our 2D imaging analysis of SEM images of the 12 selected samples (Fig. S1), unveiled a substantial ∼4-fold increase in both total lacunae density and area within the sclerotic regions compared to the trabecular regions (Fig. 7A-B). Moreover, the lacunae within the sclerotic regions exhibited significantly larger sizes and distinctive shapes compared to regular lacunae (Fig. 7C). This increase in size was primarily attributed to an elongation in the minor axis length rather than the major axis length (Fig. 7D-E), implying that lacunae in the sclerotic lesion possess greater circularity and reduced eccentricity compared to those in trabecular regions (Fig. 7F-H). Furthermore, we conducted an analysis of lacunae orientation based on our SEM images. The results revealed conspicuous differences in the distribution of angles representing the main axis of lacunae in osteosclerotic bone compared to the residual trabeculae (Fig. 8). These observations suggest that lacunae in osteosclerotic bone do not exhibit alignment with a specific orientation, which aligns with our earlier morphological observations (Fig. 5D and F). Collectively, these findings highlight that sclerotic regions of PCBMs feature a larger and irregular distribution of lacunae and pores. This observation implies an increased number of stress accumulation points in osteosclerotic bone and abnormal interaction between cells within PCBM.

**Figure 7:**
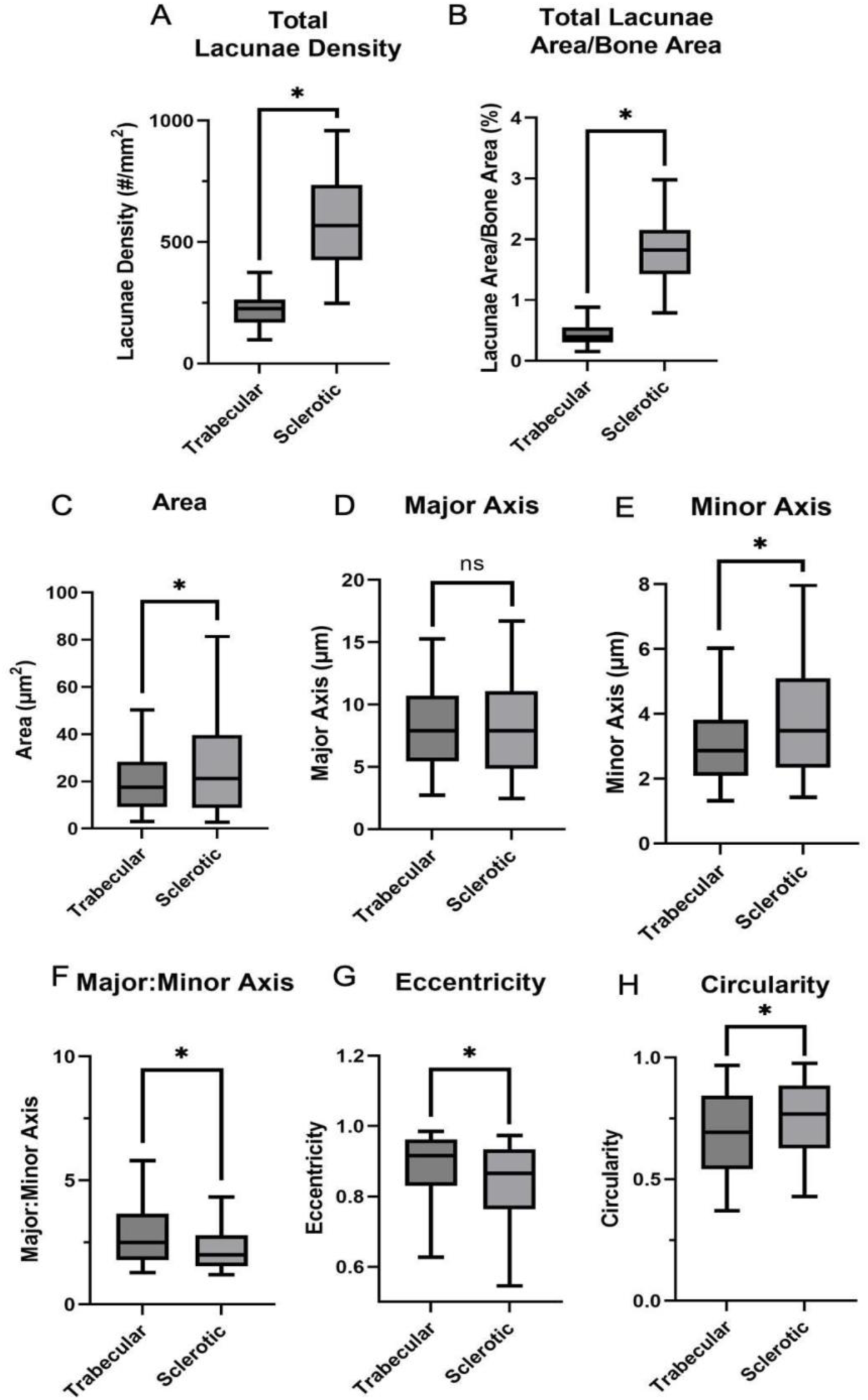
Lacunae characterization in CBM. Box plots of median +-1 and 2 Standard deviations for measurements of cadaveric vertebral PCBM samples with regions of trabecular and sclerotic bone. * p < 0.0001. n of analyzed lacunae in sclerotic areas = 3,262; n of analyzed lacunae in trabecular areas = 1,460.

**Figure 8:**
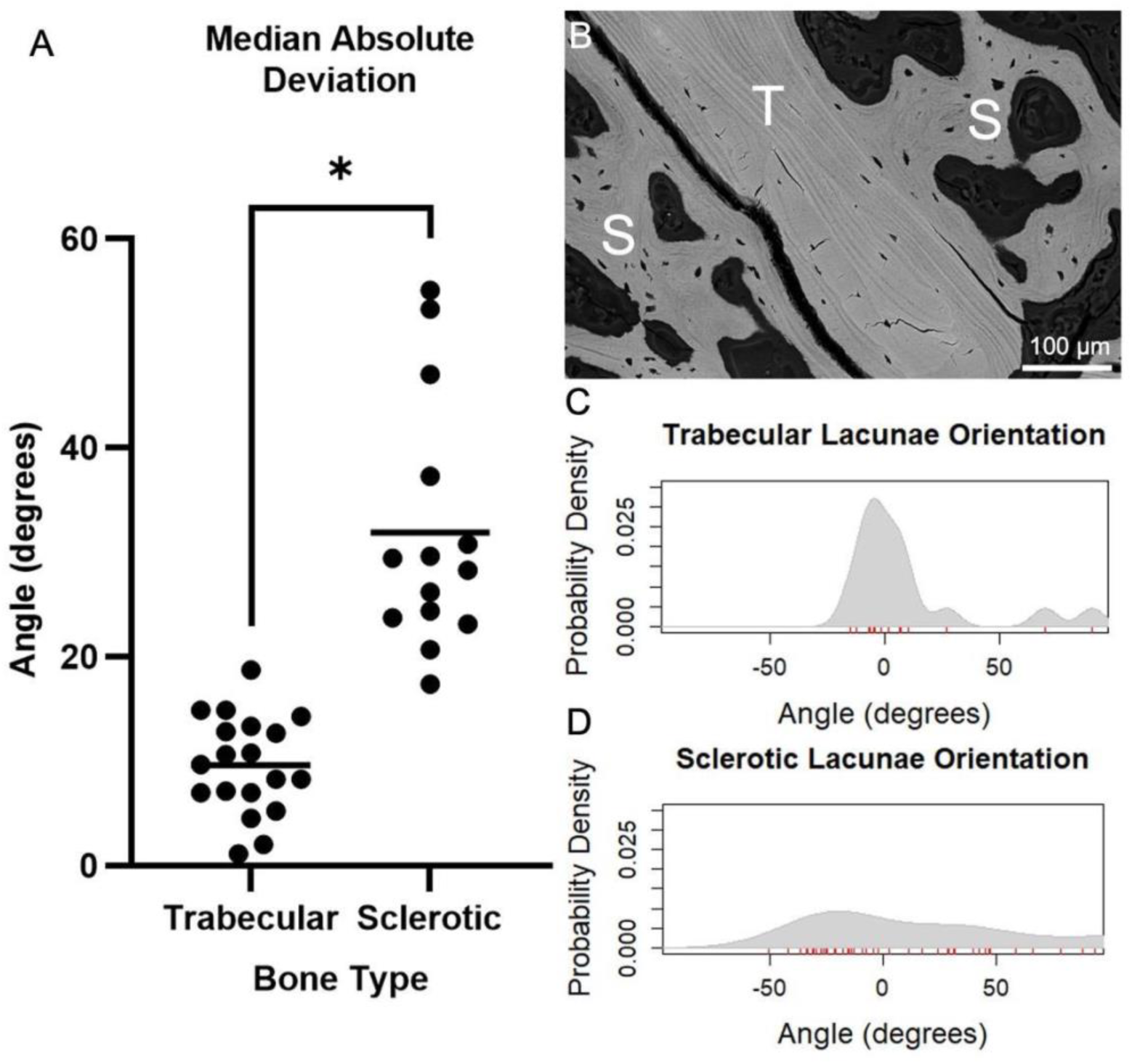
Lacunae orientation analysis in PCBM. A) Median absolute deviation of the lacunae orientation for trabecular (n=19) and sclerotic (n=14) bone regions (Total number of analyzed lacunae = 370 in trabecular bone, 718 in sclerotic bone). B) Representative quantitative back scattered electron scanning electron microscopy image of PCBM vertebrae with a trabecular region (T) and a sclerotic region (S). C) Probability density function for the lacunae orientation of trabecular bone in image B (n=16 lacunae in this image). D) Probability density function for the lacunae orientation of trabecular bone in image B in and sclerotic bone (n= 50 lacunae in this image).

### 3.7 Irregular collagen alignment in osteosclerotic bone characterize PCBM

As a result of our SEM imaging, we observed a lack of lamellar structure in the irregular PC-associated bone (Fig. 4D-F). Lamellae are critical structures in hierarchical organization of bone. They are composed of highly aligned “anisotropic” collagen fibres, which in trabecular bone are organized in lamellar packets directionally oriented following Wollf’s law (^25^), and it is the way mineralized collagen fibers provide in bone strength (^26^). Since fracture and weakness have been associated to PCBM (^27^), which lack obvious lamellae in the sclerotic PCBM regions, we performed histochemical analysis of PCBM and determine the degree of collagen alignment based on the birefringence that aligned collagen show under polarized light observation. The optical polarization properties of Sirius red allow for visualization of changes in fibrillar collagen birefringence that are related to changes in collagen isotropy (^28^). We observed strong polarization of collagen fibrils in trabecular regions, and a remarkable decreased birefringence in sclerotic regions of PCBM specimens defined as being osteosclerotic with evidence of residual trabeculae (Fig. 9A-F). Consistent with μCT measurements of the entire cores, osteolytic samples showed lower BV/TV than osteosclerotic samples with or without residual trabeculae, but here in thin sections, BV/TV of the two osteosclerotic groups was indistinguishable (Fig. S3 G). When comparing the birefringent area in these samples, we found that the highest proportion of birefringent area was in the PCBM samples defined as osteolytic (Fig S3 H). The birefringent area (as part of the total bone volume) of those defined as osteosclerotic with residual trabeculae was decreased by 20%, while that of those defined as osteosclerotic without residual trabeculae was half that of the osteolytic specimens.

**Figure 9:**
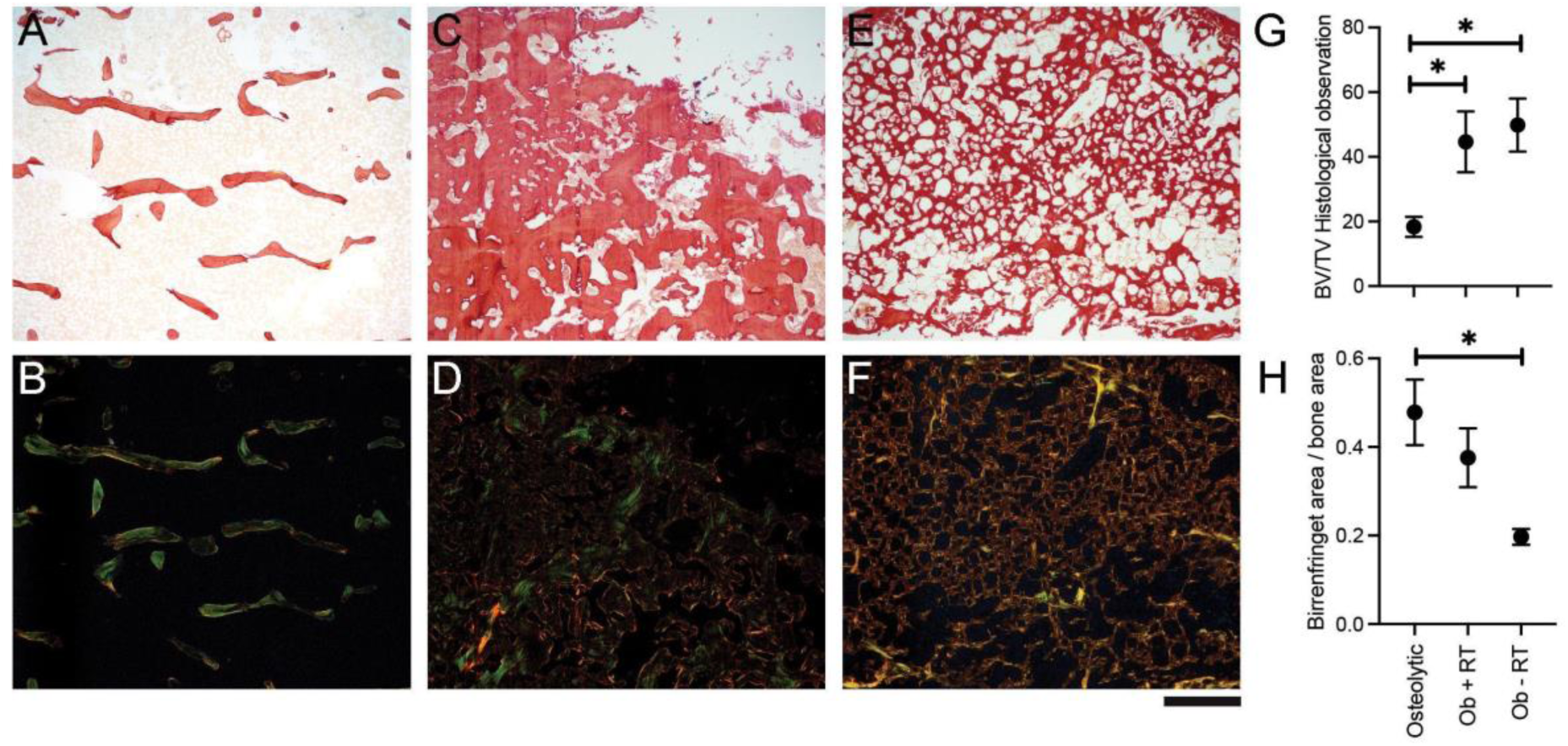
Histomorphometry of PCBM. A) Representative image of osteolytic PCBM specimen visualized with bright field. B) Polarized light imaging of A. C) Representative image of osteoblastic with residua trabeculae (OB + RT) PCBM specimen visualized in bright field. D) Polarized light imaging of C. E) Representative image of osteoblastic without residual trabeculae (OB - RT) PCBM visualized in bright field. F) Polarized light imaging of E. G) chart of mean +- - standard deviation of bone volume / total volume of different PCBM types. H) chart of birrenfringence area/ bone area of different PCBM types. * = p-value < 0.05. Staining= Picrosirius red. Scale bar = 1 mm. n = 6 osteolytic, 3 osteoblastic + residual trabeculae, 3 osteoblastic without residual trabeculae.

### 3.8 Abnormal collagen deposition is accompanied by increased proteoglycan and phosphorylated glycoprotein content in irregular bone of PCBM

Following the rationale that abnormal collagen deposition is necessarily accompanied by other ECM alterations, we went on a deeper histological characterization of PCBM. Using Goldners trichrome as a method of differentiating immature (osteoid) and mature lamellar bone (^29^), we observed strong red staining of trabecular bone, and pale blue staining of the sclerotic regions of PCBM specimens (Fig 10A). Taking advantage of the optical polarization properties of this staining method, we aligned strong birefringence in the trabecular regions defined in the bright field observations (Fig 10B). These observations independently validate our classification of predominant PCBM lesion types, and the delineation of residual trabeculae within the sclerotic lesions. When using a cationic dye such as toluidine blue, we observed that irregular PC-associated bone has a stronger affinity for this dye than regular trabecular bone (Fig. 10C), which suggests a higher negatively charged molecules content in osteosclerotic PC-associated bone. Combined, these results confirm the previous observation of irregular collagen structure and suggest the presence of other molecular alterations in the osteosclerotic PC bone.

**Figure 10:**
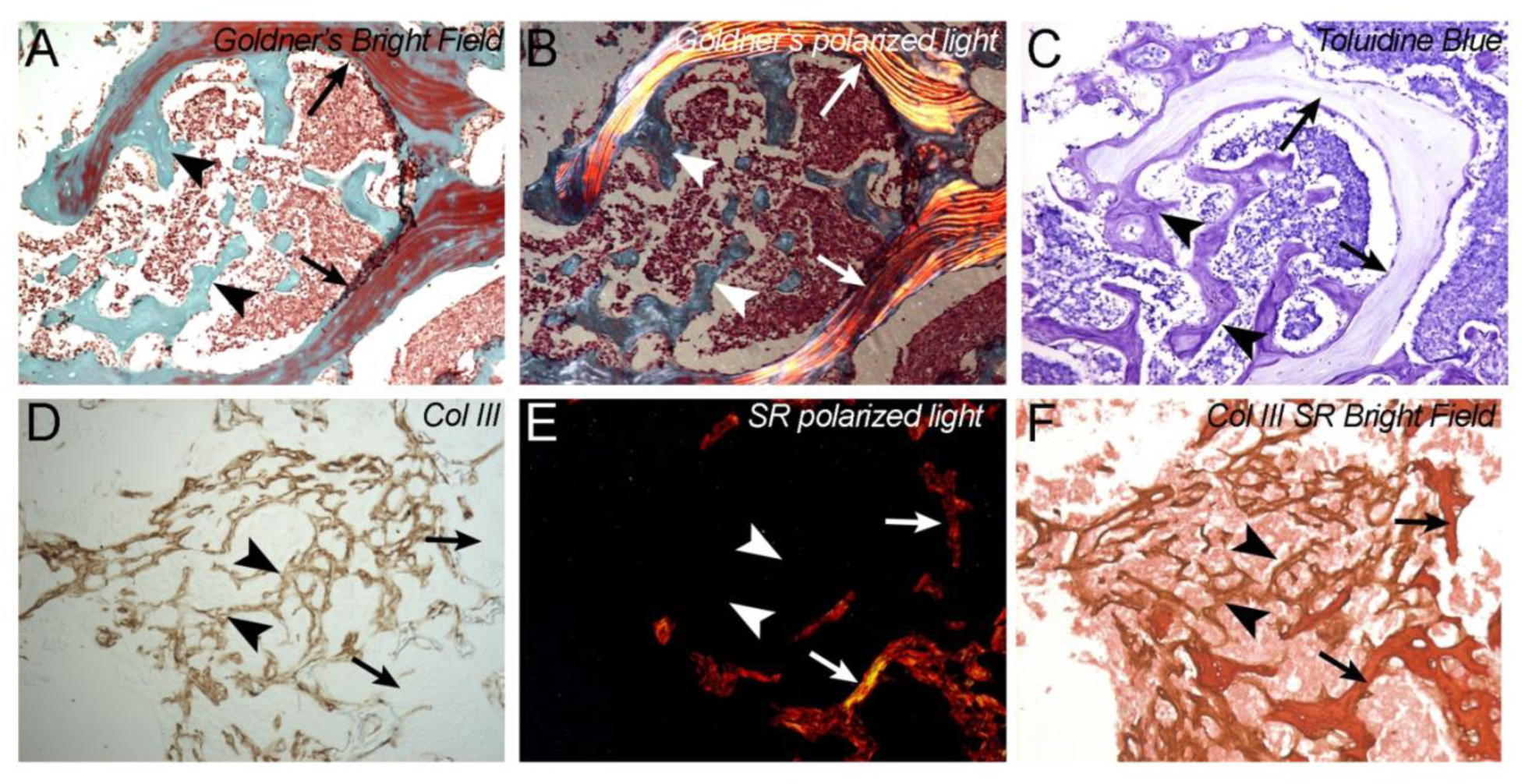
histochemistry and immunohistochemistry of extracellular matrix proteins in PCBM. A) Goldners’ trichrome staining of osteoblastic with residual trabeculae PCBM. Residual trabeculae are observed in red (arrows), while osteosclerotic bone stains light blue (Arrowheads). B) Polarized light imaging of sample slide in A. Birefringence corresponds with the red staining of residual trabeculae but is not evident in osteosclerotic bone. C) toluidine blue staining of serial section of sample in A and B, showing stronger staining of osteosclerotic bone than residual trabeculae. D) Double stain Sirius red, anti-collagen III of osteoblastic with residual trabeculae PCBM. Residual trabeculae. In bright field imaging srius red stain trabeculae (arrows) and osteosclerotic bone (arrowheads). E) polarized light imaging of D, Birrenfringence corresponds to trabeculae (bone) while osteosclerotic shows absence of birrenfringence. F) anti collagen III immunostaining of slide in D and E. Immunostaining is evident in osteosclerotic bone (arrowheads) but not in residual trabeculae (arrows

Our observations of irregular collagen organization led us to evaluate the presence of collagen III in the ECM matrix of PCBM. By using Sirius red staining on the same IHC stained slides we could simultaneously observe collagen alignment and the presence of collagen III. We observed that PCBM exhibit areas that strongly stain with collagen III immunostaining forming an irregular network in the intertrabecular space (Fig 10D). While Sirius red staining shows absence of birefringence of these collagen III-rich areas, but strong birefringence collagen III negative trabeculae (Fig. 10E). These results demonstrate the presence of collagen III as a major component of sclerotic PCBM.

Because of these observed differences we expanded the immunohistochemical analysis of PCBM to a group of bone proteins. We analyzed collagen I (Col I), the most abundant structural protein of bone, and observed increased staining for Col I in osteosclerotic bone compared to residual trabecula and showed enhanced staining in the osteosclerotic areas (Fig. 11A).

**Figure 11:**
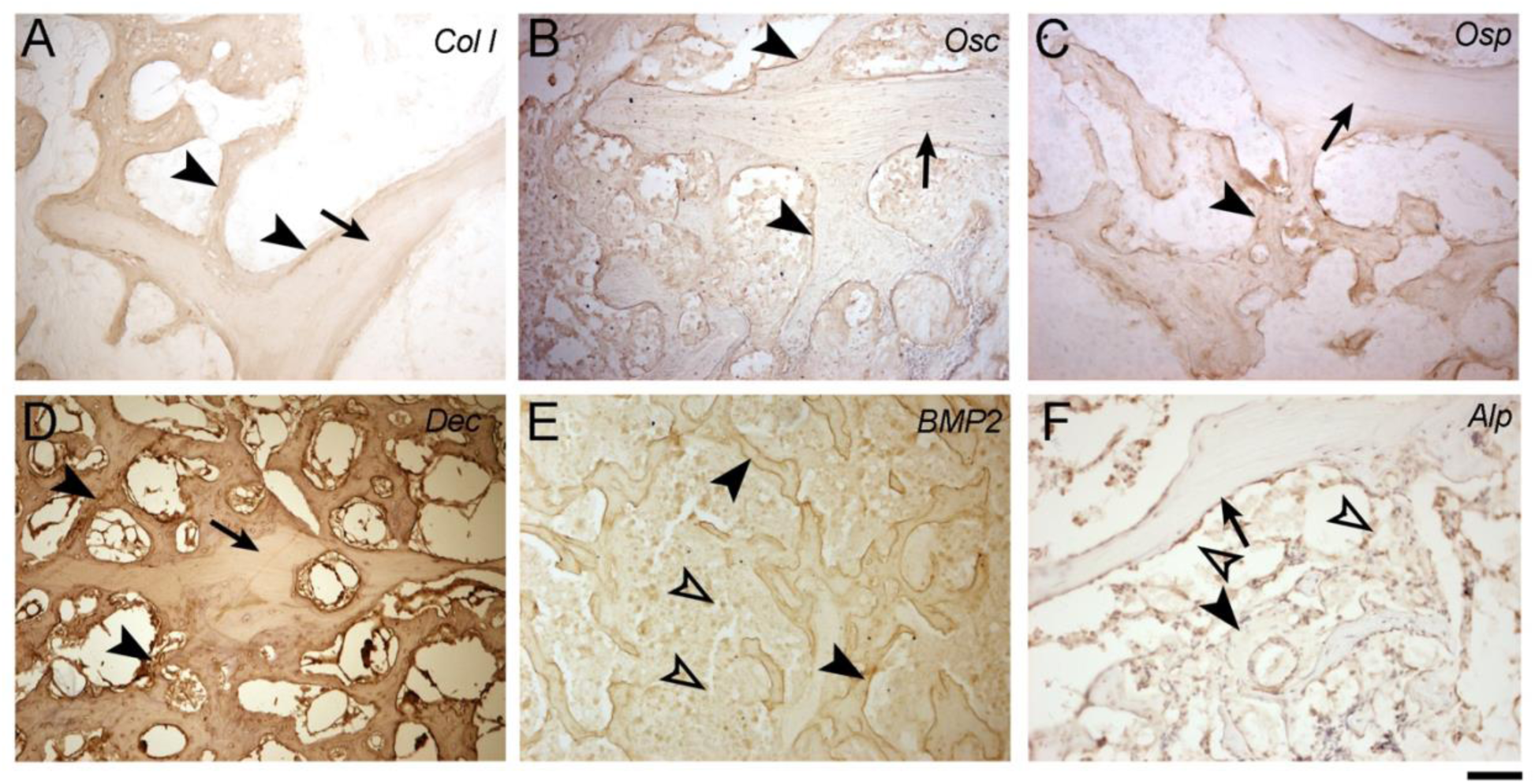
histochemistry and immunohistochemistry of extracellular matrix proteins in PCBM. A) Immuhistochemical staining for collagen I, showing stronger signal in osteosclerotic bone (arrowheads) than in residual trabeculae that are also positive (arrows). B) Immuhistochemical staining for osteocalcin mild staining in the matrix of trabecular (arrow) and sclerotic (arrowheads)bone. Intersticcial staining in non-mineralized tissue s also obseved. C) Immuhistochemical staining for osteopontin showing stronger signal in osteosclerotic bone (arrowheads) than in residual trabeculae (arrows). D) Immunostaining for decorin shows stronger immunostaining in sclerotic bone (arrowheads) than in residual trabeculae (arrow). E) Immuhistochemical staining for bone morphogenetic protein 2. Strong immunostaining is evident on the bone surfaces (arrows) but is also positive in interstitial cells (open arrowheads). F) Immuhistochemical staining for alkaline phosphatase. The immunostaining is minimum in pre-existing trabeculae (arrow) and osteosclerotic bone (arrowheads), but strong in cells localized in the bone surfaces (osteoblasts, open arrowheads).

Osteocalcin (Osc), the most abundant non-collagenous protein in bone, showed a mild staining in the matrix of osteosclerotic and trabecular bone, but a marked staining in bone surfaces and cells (Fig. 11B). We detected a higher immunostaining in osteosclerotic bone for Osteopontin (Osp), a highly phosphorylated protein from the SIBLING family (Fig. 11C). The increased affinity for cationic dyes suggests that polyanionic molecules are highly expressed in the irregular PC-associated bone. Proteoglycans are the trademark of polyanionic molecules in the extracellular matrix. Decorin, is critical for both collagen fiber organization and biomineralization (^30, 31^). Through IHC we observed stronger staining for decorin (Dec) in the extracellular matrix of osteosclerotic than lamellar bone (Fig. 11D). BMP2 instead showed minimum staining in the matrix, but intense staining in the surface of trabecular or sclerotic bone, as well as in the cells in the intertrabecular space (Fig. 11E). The formation of calcium crystals in bone is highly dependent on Ca^2+^ and PO_4_ concentration, which in mature bone is generated by the presence of alkaline phosphatase, an enzyme that catalyzed the release of PO4 from different substrates (^32^), while in cartilage-based osteogenesis and in woven bone, the mineralization of cartilage matrix is mediated by matrix vesicles, membranous structures that generate a microenvironment capable of crystal precipitation (^33^). To observe which mechanism of mineralization could be predominant in irregular PC-associated bone, we performed IHC for alkaline phosphatase and PHOSPHO-1 a well-accepted marker for matrix vesicles. Alkaline phosphatase (ALP) showed no staining in the bone matrix but abundant staining in osteoblasts and osteocytes in both lamellar and osteosclerotic bone (Fig. 11F). As expected, we also observed elevated ALP in the cellular component of the tissues due to the known expression of ALP in metastatic PC cells (^34^). We observed no staining for cartilage markers Col 2 and Aggrecan, and no evidence of matrix vesicle mineralization marker PHOSPHO1 (Fig. S4). Combined these results confirm the altered structure of the bone matrix in irregular PC-associated bone, with no cartilage switch but resembling a reparative bone matrix, with increased collagen I and III and increased mineralization-related proteins and proteoglycans.

## 4. DISCUSSION

By generating tremendous intractable pain and increasing the risk of fractures, BM are one of the most severe consequences of PC. Contradictorily, PCBM are osteoblastic lesions with increased mineral content, but the mechanisms of pain and fracture, as well as the role of PC cells in this process are not clearly established. In this study, we conducted an extensive exploration of the structural and mechanical characteristics of PCBM, comprising the largest sample number reported to date, we complemented our study with histological analysis and a preliminary immunohistochemical characterization of the PCBM extracellular matrix. Our findings reveal that structural disparities in PCBM, marked by the emergence of sclerotic PC-associated bone, are characterized by altered bone distribution, abnormal collagen organization, and increased porosity. These structural changes may ultimately underlie the bone’s weakness and the heightened fracture risk observed in prostate cancer patients. Furthermore, our analysis indicates that these alterations in PCBM are not attributed to differences in the calcium content, or nano-scale hardness or elastic modulus. Instead, they appear to arise from a loss of three-dimensional organization of the collagen matrix. In essence, our research sheds light on the complex interplay of structural and mechanical factors in PCBM, providing valuable insights into the mechanisms behind pain and fractures in prostate cancer patients.

Early x-ray imaging showed that PCBM could generate sclerotic lesions with increased radiopacity, radio lucid lesions and lesions that combine these two phenotypes described as mixed. These three phenotypes have been largely described; however, these descriptions have been mostly based on patients’ x-ray, or murine bone analysis after PC cells inoculation (^35, 36^). High resolution studies on human samples have been sparse, and with limited sample number. One high-resolution observation of human vertebrae with PCBM used a synchrotron-energized mCT scanner, they analyzed 7 mm isometric cubic sections of spine (n=15) from a single patient who died of PC, with a pixel size of 23.2 um and a step size of 18.56 um reported increased bone surface and trabecular number in the PCBM compared to normal vertebrae, but no difference in trabecular thickness (^9^). The same group performed a higher resolution analysis at 6 um isometric voxel size, of 4 mm side cubes from vertebrae with PCBM obtained from two patients, and described a higher BV/TV, bone surface, trabecular number, trabecular connectivity, anisotropy and fractal dimensions in PCBM (^10^). More recently, mCT scanning at an isometric voxel size of 10.5 μm was used to analyzed 5 mm diameter cores obtained from cadaveric vertebra with metastasis from two men who died of PC, 3 patients who died of breast cancer and 3 from lung cancer (^11^). By complimenting this morphological analysis with mechanical testing and biochemical analysis, the PCBM cores were scored as sclerotic, lytic, or mixed, but for the statistical analysis, all samples were grouped according to imaging classification, thus no conclusion can be made from the specific cancer type. Finally, a μCT analysis of metastatic vertebrae, performed at an isometric voxel size of 24.5 um describe PCBM as osteoblastic, lytic and/or mixed lesions on samples obtained from two patients (^12^).

By comparison, our observations, in the largest cohort of PCBM described so far, and non-PC controls, find agreement with these previous descriptions. We found increased BV/TV, trabecular number, and connectivity in samples with osteoblastic PCBM, and no increase in trabecular thickness, and we also observed a variety of patterns from sclerotic to lytic lesions. Notably, our analysis led to a distinct description of osteoblastic samples, based on the pre-existing osteolytic activity in osteoblastic samples observed by the absence of remaining trabeculae. This differential pattern was previously described in the histological observation of the same sample cohort (^8^). They described osteoblastic lesions as “…consisted mostly of woven bone, with small amounts of osteoid and inclusions of native bone trabeculae appearing as lamellar bone… …In some cases an intact lamellar trabecular network remained, with woven bone around the edges of the normal bone.”. Our work, benefiting from volumetric and three-dimensional analysis using high-resolution μCT scans, unveils these distinct patterns, suggesting the involvement of two different mechanisms, or timing, of osteolytic and osteoblastic activity. These processes need further study since different vertebral lesions can present as osteolytic and osteoblastic in a given patient.

Because of its morphological characteristic, sclerotic PCBM lesions are generally described as “woven bone” (^37–39^). Woven bone is defined as a temporary mineralized bone matrix present during the initial synthesis of intramembranous lamellar bone (^40^), in both embryonic development and fracture repair (^18^), while its relationship with trabecular bone is still not clear. With a collagenous matrix composed of fibrils between 0.1 and 3um thickness randomly oriented, the calcium crystals also show a random orientation generating a highly mineralized matrix but porous at the micron level (^41^). Woven bone is transient. After its advantageous fast synthesis and mineralization, it is remodeled as lamellar bone, in a process that starts by deposition of bone lamellae in the surface of woven bone (^18, 42^). Woven bone is histologically characterized by the presence of a “meshwork of collagen bundles which bear no fixed relationship either to the lacunae or vascular spaces of the tissue” (^40^). Woven bone lacks lamellar structure, and is characterized as having rounded lacunae within a fibrillar collagen I matrix (^43^) with some differences in protein composition (^44^).

Our observations of sclerotic PCBMs partially agree with these descriptions. We observed irregular matrix deposition, with poor collagen alignment and rounded lacunae, which in turn are not aligned with any structures, and lack cartilage proteins. However, we did not observe any lamellar or aligned collagen deposition on the surface of woven bone, and no resorption of this structure that could lead to a later stage of woven bone life cycle (^18^). Our observation of collagen III accumulation in sclerotic PCBM matrix is not traditionally described in woven bone. It is, however, described during the bone wound healing process (^45^), harkening Dvorak’s description of cancer as “wounds that do not heal” (^46^). What cells are responsible for collagen III deposition and how this matrix mineralizes needs to be investigated, but our preliminary observations suggests that, unlike woven bone, PCBM mineralize by a matrix vesicles-independent process, and perhaps directed by the presence of SIBLING proteins such as osteopontin, and proteoglycans, such as decorin, which are known to play key roles in fibrilogenesis and biomineralization.

Bone hierarchical structure is critical for its mechanical properties. The organization of collagen fibers and lamellar structure allows elastic and inelastic deformation which enhance the material toughness (^47^). Additionally, lamellae are instrumental in resisting crack initiation and propagation (^23^). These lamellar structures counteract the “weakness” that results from the stress accumulation in the pores of biomineralized composite materials (^48^). Furthermore, the small canaliculi are initiation points for microcrack that allows inelastic deformation, increasing bone toughness, at the time that a dense collagen network prevents crack propagation, preventing bone fracture (^24^). The structural alterations that we observed in PCBM, theoretically would lead to increased weakness. Due to the enhanced porosity caused by a larger lacunae number, size, and irregular shape, the chances of crack initiation would increase under tensile and compressive forces, while the absence of trabecular structure would not prevent crack propagation. Interfaces between two different materials are also stress accumulation areas, as we observed in our samples, these material differences may also be a starting point of cracks. With minimum or non-bonding mechanisms between irregular PCBM associated bone and pre-existing trabeculae, these cracks may easily propagate through the interface, further increasing the risk of fracture.

The alteration of collagenous and non-collagenous ECM of irregular PC-associated bone that we describe in this article are novel in human tissues, but some interesting observations have been explored previously in a variety of animal models. High Performance Liquid Chromatography and Raman spectroscopy analysis of osteoblastic murine bone lesions generated after HELA cell injection exhibited lower collagen crosslinking and mineral crystallinity, suggesting an alteration in the mineralization process of bone (^49^). Concordantly, Raman spectroscopy analysis of LNCaP C4-2B inter-tibial osteoblastic lesions were characterized by the presence of collagen with a lower mineral ratio than normal bone, and the calcium component showed a lower crystallinity and higher carbonate substitution, compared to the non-affected contralateral tibiae (^35^). A similar analysis of sclerotic lesions caused by inoculation of PCa-2b cells revealed increased bone volume, bone volume fraction, bone mineral content and mineral density compared to control (^36^). By quantitative polarized microscopy the authors determined a reduced collagen alignment in the newly formed bone, and by using x-ray-diffraction they determined a lower crystal alignment in the PC-induced bone. Our results on human samples find concordance with some of these results, in terms of collagen arrangement, and further analysis in the mineral phase are needed to establish definitive conclusions.

## ACKNOWLEDGMENTS

The authors thank the patients and their families.

