## Supplement for "Sclerotic prostate cancer bone metastasis: woven bone lesions with a twist"

Running title: Structure of prostate cancer bone metastasis

### Table S1

Demographics and clinical data of patients

| Patient | Vertebrae | SampNo | Race | Glessons primary | PSA at Ddx | Final Serum PSA (ng/mL) | Age at Diagnosis | D PSA | Age @ death (years) | survival from dx (years) |
|---|---|---|---|---|---|---|---|---|---|---|
| 1 | L1 | 42 | C | 4+3 | 10.5 | 21.16 | 48.08 | 37.58 | 52.15 | 4.07 |
| | L2 | 72 | | | | | | | | |
| | L3 | 73 | | | | | | | | |
| 2 | L4 | 43 | C | 4+5 | 2.6 | 0.5 | 52.73 | 50.13 | 58.06 | 5.33 |
| | L5 | 60 | | | | | | | | |
| 3 | L2 | 16 | M | 5+4 | 105 | 620.5 | 57.84 | -47.16 | 59.94 | 2.09 |
| 4 | L1 | 47 | C | 4+4 | 6.2 | 1 | 56.54 | 50.34 | 60.16 | 3.62 |
| | L2 | 50 | | | | | | | | |
| | L3 | 56 | | | | | | | | |
| 5 | L1 | 21 | C | 5+4 | 76 | 254.49 | 59.70 | -16.30 | 61.00 | 1.30 |
| | T11 | 28 | | | | | | | | |
| 6 | L1 | 43 | C | 4+5 | 6 | 0.15 | 59.22 | 53.22 | 61.60 | 2.38 |
| | L2 | 58 | | | | | | | | |
| 7 | L2 | 62 | C | 4+3 | 20.6 | 967.48 | 50.68 | 30.08 | 62.22 | 11.54 |
| | L4 | 66 | | | | | | | | |
| | L1 | 74 | | | | | | | | |
| 8 | L5 | 13 | C | 4+5 | 15.8 | 2534.86 | 59.17 | 43.37 | 63.44 | 4.27 |
| 9 | T11 | 23 | C | 4+3 | 64.6 | 3869 | 59.58 | -5.02 | 64.04 | 4.46 |
| 10 | L4 | 10 | C | 3+5 | 8.6 | 106.1 | 63.12 | 54.52 | 64.71 | 1.59 |
| | L3 | 15 | | | | | | | | |
| | L5 | 30 | | | | | | | | |
| 11 | L5 | 6 | C | 4+5 | 1000.1 | 767.6 | 61.99 | -938.11 | 65.19 | 3.19 |
| | L4 | 11 | | | | | | | | |
| | L2 | 17 | | | | | | | | |
| 12 | L2 | 4 | M | 4+3 | 4 | 380.95 | 61.56 | 57.56 | 66.43 | 4.87 |
| | L1 | 14 | | | | | | | | |
| | L3 | 20 | | | | | | | | |
| 13 | L2 | 52 | C | 4+5 | 1.7 | 58.74 | 63.38 | 61.68 | 67.20 | 3.82 |
| 14 | L2 | 68 | C | 4+5 | 0 | 0.72 | 58.49 | 58.49 | 67.28 | 8.79 |
| 15 | L3 | 63 | C | 4+5 | 14 | 454 | 64.97 | 50.97 | 67.49 | 2.52 |
| | L1 | 64 | | | | | | | | |
| 16 | L1 | 54 | M | 4+4 | 226 | 328.3 | 66.09 | -159.91 | 69.16 | 3.07 |
| 17 | L5 | 69 | C | 3+4 | 13 | 104.4 | 62.67 | 49.67 | 70.12 | 7.45 |
| 18 | L4 | 49 | C | 4+4 | 20 | 90.94 | 63.49 | 43.49 | 71.21 | 7.72 |
| 19 | T12 | 26 | C | 7 | 33.9 | 105.6 | 62.07 | 28.17 | 71.64 | 9.56 |
| 20 | L5 | 36 | C | 9 | | 119.7 | 61.50 | 61.50 | 72.39 | 10.89 |
| | L4 | 39 | | | | | | | | |
| | L1 | 57 | | | | | | | | |
| 21 | L5 | 33 | C | 4+3 | 12 | 2511.73 | 63.66 | 51.66 | 72.65 | 8.99 |
| 22 | L4 | 41 | C | 3+4 | 5.4 | 2816 | 59.65 | 54.25 | 72.88 | 13.23 |
| 23 | L5 | 38 | M | 4+4 | 54.8 | 290.1 | 70.09 | 15.29 | 72.93 | 2.84 |
| | L2 | 61 | | | | | | | | |
| | L4 | 71 | | | | | | | | |
| 24 | L3 | 29 | C | 7 | 42 | 3.3 | 61.68 | 19.68 | 73.14 | 11.46 |
| | L5 | 51 | | | | | | | | |
| 25 | L2 | 22 | C | 8 | 41 | 359 | 62.84 | 21.84 | 75.43 | 12.59 |
| | L4 | 65 | | | | | | | | |
| | L3 | 70 | | | | | | | | |
| 26 | L4 | 5 | C | 3+4 | 13.4 | 547 | 70.36 | 56.96 | 76.02 | 5.66 |
| | L2 | 7 | | | | | | | | |
| | L1 | 24 | | | | | | | | |
| 27 | L1 | 25 | C | 4+3 | 12 | 3700 | 59.84 | 47.84 | 76.59 | 16.75 |
| | L4 | 3 | | | | | | | | |
| | L3 | 8 | | | | | | | | |
| 28 | L1 | 2 | C | 3+3 | 10 | 73.1 | 65.15 | 55.15 | 76.84 | 11.69 |
| | L4 | 27 | | | | | | | | |
| 29 | L4 | 44 | M | 5+5 | 12 | 762.3 | 72.57 | 60.57 | 77.13 | 4.56 |
| | L3 | 48 | | | | | | | | |
| 30 | L1 | 34 | C | 4+4 | 10.2 | 1250.4 | 63.22 | 53.02 | 79.52 | 16.31 |
| 30 | L5 | 67 | C | 4+5 | 49.1 | 5.63 | 78.36 | 29.26 | 80.83 | 2.48 |
| 32 | T11 | 35 | C | 5+5 | 27.2 | 16.6 | 75.44 | 48.24 | 81.02 | 5.58 |
| 33 | L3 | 19 | M | 3+3 | 6.2 | 2295 | 69.93 | 63.73 | 81.71 | 11.78 |
| 34 | L3 | 45 | C | unk | | 63.23 | 68.63 | 68.63 | 82.53 | 13.89 |
| 35 | L5 | 55 | C | 2+3 | 0 | 30.7 | 68.49 | 68.49 | 82.98 | 14.49 |
| 36 | L2 | 40 | C | 3+4 | | 114.83 | 72.98 | 72.98 | 83.47 | 10.49 |
| 37 | L1 | 31 | C | 3+4 | 43 | 240.65 | 74.62 | 31.62 | 83.70 | 9.08 |
| 38 | L1 | 59 | C | 4+3 | 12.4 | 721.9 | 73.77 | 61.37 | 85.67 | 11.90 |
| | L3 | 75 | | | | | | | | |
| 39 | T12 | 32 | M | 4+3 | 10 | 1508.4 | 74.30 | 64.30 | 86.06 | 11.76 |
| 40 | L4 | 9 | C | UNK | | 413.2 | 65.98 | 65.98 | 86.23 | 20.25 |
| | L3 | 12 | | | | | | | | |
| | L5 | 76 | | | | | | | | |
| 41 | L4 | 18 | C | 3+5 | 11.1 | 349.05 | 72.49 | 61.39 | 90+ | 20.21 |
| 42 | L4 | 25 | C | unk | 0 | 1455.9 | 78.65 | 78.65 | 90+ | 11.18 |
| | L2 | 37 | | | | | | | | |

Table S2

Antibodies used in IHC.

| Primary antibodies | | | |
|---|---|---|---|
| Antigen | Host species | Brand | Cat No |
| Aggrecan | Rb | Abcam | 36861 |
| Col 3 | Ms | Abcam | 6310 |
| Alp | Rb | Abcam | 224335 |
| Col 1 | Rb | Bioss | BS-0578R |
| Osc | Rb | Abcam | 93876 |
| PHOSPHO1 | Rb | Abcam | 272655 |
| BMP2 | Rb | LSBio | B12549 |
| Dec | Rb | Millipore | ABT276 |
| Osn | Rb | Immundiagnostik | A4225 |
| BSP | Ms | Immundiagnostik | A4232 |
| Col2 | Ms | Millipore | MAB8887 |
| Secondary antibodies - HRP conjugated | | | |
| Mouse Igg | Goat | Abcam | 205719 |
| Rabbit Igg | Goat | Abcam | 205718 |

### Table S3 uCT values of PCBM

| Samp No | Patient | BV/TV | Conn-Dens. | Tb.N | Tb.Th | Tb.Sp | Tb.Th | Tb.Sp | BMD |
|---|---|---|---|---|---|---|---|---|---|
| 1 | 27 | 0.791 | 193.4 | 7.7 | 0.242 | 0.119 | 0.106 | 0.116 | 910.5 |
| 2 | 28 | 0.788 | 417.7 | 11.8 | 0.131 | 0.057 | 0.061 | 0.029 | 910.5 |
| 3 | 27 | 0.760 | 65.6 | 6.5 | 0.252 | 0.165 | 0.100 | 0.117 | 939.5 |
| 4 | 12 | 0.732 | 139.4 | 8.3 | 0.168 | 0.119 | 0.064 | 0.083 | 870.9 |
| 5 | 26 | 0.723 | 178.5 | 9.5 | 0.153 | 0.121 | 0.058 | 0.094 | 901.0 |
| 6 | 11 | 0.698 | 500.4 | 10.6 | 0.136 | 0.099 | 0.066 | 0.074 | 883.2 |
| 7 | 26 | 0.682 | 478.9 | 10.2 | 0.134 | 0.105 | 0.064 | 0.081 | 864.1 |
| 8 | 27 | 0.663 | 218.2 | 8.7 | 0.174 | 0.126 | 0.086 | 0.085 | 902.6 |
| 9 | 42 | 0.622 | 280.8 | 10.4 | 0.109 | 0.093 | 0.045 | 0.051 | 891.6 |
| 10 | 10 | 0.621 | 147.5 | 7.4 | 0.135 | 0.182 | 0.049 | 0.158 | 871.7 |
| 11 | 11 | 0.616 | 95.0 | 6.0 | 0.185 | 0.167 | 0.084 | 0.093 | 879.9 |
| 12 | 42 | 0.600 | 302.9 | 10.5 | 0.106 | 0.097 | 0.043 | 0.052 | 869.4 |
| 13 | 8 | 0.578 | 378.5 | 8.0 | 0.117 | 0.170 | 0.047 | 0.125 | 855.6 |
| 14 | 12 | 0.551 | 270.0 | 8.1 | 0.120 | 0.130 | 0.060 | 0.065 | 848.7 |
| 15 | 10 | 0.548 | 130.1 | 5.1 | 0.143 | 0.294 | 0.057 | 0.228 | 880.9 |
| 16 | 3 | 0.528 | 627.1 | 9.4 | 0.105 | 0.099 | 0.056 | 0.064 | 842.5 |
| 17 | 11 | 0.513 | 143.8 | 6.3 | 0.153 | 0.195 | 0.069 | 0.111 | 843.0 |
| 18 | 44 | 0.500 | 74.5 | 4.3 | 0.182 | 0.319 | 0.092 | 0.192 | 856.1 |
| 19 | 35 | 0.491 | 59.8 | 3.8 | 0.174 | 0.368 | 0.074 | 0.285 | 843.5 |
| 20 | 12 | 0.479 | 282.6 | 8.1 | 0.101 | 0.131 | 0.043 | 0.065 | 819.2 |
| 21 | 5 | 0.441 | 153.7 | 5.7 | 0.127 | 0.207 | 0.059 | 0.177 | 839.6 |
| 22 | 25 | 0.416 | 34.5 | 3.8 | 0.182 | 0.371 | 0.083 | 0.231 | 836.7 |
| 23 | 9 | 0.415 | 374.4 | 6.0 | 0.078 | 0.199 | 0.033 | 0.153 | 852.4 |
| 24 | 26 | 0.392 | 146.3 | 4.6 | 0.114 | 0.266 | 0.047 | 0.196 | 821.5 |
| 25 | 34 | 0.200 | 9.6 | 1.7 | 0.110 | 0.603 | 0.039 | 0.199 | 834.3 |
| 25 | 45 | 0.380 | 332.6 | 6.8 | 0.106 | 0.157 | 0.057 | 0.085 | 862.3 |
| 26 | 19 | 0.341 | 98.6 | 3.7 | 0.140 | 0.314 | 0.079 | 0.224 | 862.6 |
| 27 | 28 | 0.316 | 497.7 | 4.5 | 0.073 | 0.258 | 0.041 | 0.232 | 820.5 |
| 28 | 5 | 0.301 | 170.5 | 3.6 | 0.132 | 0.330 | 0.068 | 0.202 | 849.2 |
| 29 | 24 | 0.297 | 55.3 | 2.3 | 0.146 | 0.517 | 0.063 | 0.296 | 831.3 |
| 30 | 10 | 0.296 | 60.7 | 2.2 | 0.139 | 0.540 | 0.065 | 0.334 | 858.2 |
| 31 | 39 | 0.282 | 110.1 | 3.5 | 0.122 | 0.323 | 0.064 | 0.270 | 808.5 |
| 32 | 41 | 0.240 | 106.3 | 2.5 | 0.113 | 0.458 | 0.060 | 0.270 | 785.2 |
| 33 | 21 | 0.208 | 16.7 | 1.9 | 0.142 | 0.583 | 0.076 | 0.210 | 852.0 |
| 34 | 31 | 0.205 | 40.1 | 1.9 | 0.141 | 0.574 | 0.077 | 0.296 | 827.9 |
| 36 | 20 | 0.195 | 51.6 | 3.0 | 0.120 | 0.338 | 0.072 | 0.220 | 841.6 |
| 37 | 45 | 0.189 | 109.5 | 3.2 | 0.093 | 0.334 | 0.058 | 0.212 | 812.6 |
| 38 | 23 | 0.183 | 10.2 | 1.2 | 0.181 | 0.899 | 0.083 | 0.551 | 871.4 |
| 39 | 20 | 0.173 | 38.2 | 2.0 | 0.108 | 0.526 | 0.047 | 0.268 | 806.5 |
| 40 | 38 | 0.172 | 43.1 | 1.2 | 0.154 | 0.955 | 0.070 | 0.442 | 808.6 |
| 41 | 22 | 0.167 | 7.5 | 1.3 | 0.169 | 0.823 | 0.064 | 0.330 | 877.4 |
| 42 | 1 | 0.161 | 32.6 | 1.9 | 0.113 | 0.555 | 0.060 | 0.288 | 801.7 |
| 43 | 6 | 0.160 | 24.6 | 1.7 | 0.110 | 0.605 | 0.042 | 0.255 | 818.6 |
| 44 | 29 | 0.156 | 44.5 | 1.7 | 0.132 | 0.617 | 0.070 | 0.349 | 846.5 |
| 46 | 37 | 0.146 | 102.8 | 2.3 | 0.085 | 0.461 | 0.043 | 0.260 | 804.9 |
| 47 | 4 | 0.138 | 4.0 | 1.0 | 0.180 | 1.046 | 0.079 | 0.375 | 840.6 |
| 48 | 29 | 0.135 | 39.9 | 1.7 | 0.110 | 0.643 | 0.063 | 0.359 | 813.1 |
| 49 | 18 | 0.133 | 57.3 | 2.0 | 0.095 | 0.529 | 0.046 | 0.366 | 819.8 |
| 50 | 4 | 0.132 | 5.1 | 1.3 | 0.139 | 0.807 | 0.051 | 0.212 | 850.0 |
| 51 | 24 | 0.128 | 48.3 | 1.3 | 0.102 | 0.796 | 0.062 | 0.437 | 844.1 |
| 52 | 13 | 0.127 | 14.2 | 1.5 | 0.112 | 0.700 | 0.050 | 0.243 | 833.5 |
| 53 | 2 | 0.124 | 39.3 | 1.8 | 0.097 | 0.569 | 0.044 | 0.271 | 814.9 |
| 54 | 16 | 0.119 | 16.4 | 1.6 | 0.103 | 0.623 | 0.047 | 0.200 | 875.0 |
| 55 | 36 | 0.118 | 15.7 | 1.2 | 0.126 | 0.912 | 0.058 | 0.475 | 850.9 |
| 56 | 4 | 0.114 | 5.2 | 1.2 | 0.125 | 0.855 | 0.044 | 0.257 | 838.5 |
| 57 | 20 | 0.113 | 7.4 | 1.3 | 0.102 | 0.782 | 0.034 | 0.340 | 844.8 |
| 58 | 6 | 0.110 | 13.6 | 1.7 | 0.097 | 0.600 | 0.036 | 0.269 | 821.5 |
| 59 | 40 | 0.110 | 6.4 | 1.2 | 0.157 | 0.841 | 0.078 | 0.353 | 881.9 |
| 60 | 2 | 0.109 | 20.2 | 1.8 | 0.097 | 0.579 | 0.042 | 0.310 | 821.2 |
| 61 | 23 | 0.108 | 4.2 | 1.1 | 0.174 | 0.947 | 0.093 | 0.305 | 876.3 |
| 62 | 7 | 0.095 | 4.8 | 1.2 | 0.125 | 0.853 | 0.053 | 0.249 | 830.8 |
| 63 | 15 | 0.091 | 29.6 | 1.3 | 0.104 | 0.787 | 0.053 | 0.402 | 825.3 |
| 64 | 15 | 0.090 | 4.1 | 0.9 | 0.135 | 1.104 | 0.063 | 0.388 | 863.4 |
| 65 | 25 | 0.088 | 7.2 | 1.1 | 0.098 | 0.924 | 0.039 | 0.321 | 861.0 |
| 66 | 7 | 0.088 | 3.8 | 1.0 | 0.132 | 0.964 | 0.061 | 0.279 | 835.0 |
| 67 | 32 | 0.087 | 5.3 | 1.2 | 0.120 | 0.813 | 0.056 | 0.403 | 837.5 |
| 68 | 14 | 0.084 | 10.8 | 1.0 | 0.095 | 1.010 | 0.037 | 0.300 | 828.7 |
| 69 | 17 | 0.083 | 2.4 | 0.9 | 0.117 | 1.145 | 0.040 | 0.337 | 855.5 |
| 70 | 25 | 0.080 | 4.5 | 1.0 | 0.112 | 1.009 | 0.053 | 0.356 | 864.8 |
| 71 | 23 | 0.080 | 3.2 | 1.0 | 0.129 | 0.975 | 0.054 | 0.269 | 872.0 |
| 72 | 1 | 0.079 | 22.1 | 1.3 | 0.069 | 0.790 | 0.025 | 0.259 | 807.1 |
| 73 | 1 | 0.078 | 7.4 | 1.1 | 0.083 | 0.902 | 0.029 | 0.340 | 849.0 |
| 74 | 7 | 0.074 | 4.8 | 1.0 | 0.101 | 0.981 | 0.041 | 0.335 | 816.1 |
| 75 | 40 | 0.073 | 2.9 | 1.0 | 0.135 | 1.017 | 0.048 | 0.369 | 885.9 |

Table S4, micro-CT values of age matched control samples

| Sample No | BV/TV | Tb.Sp | Tb.N | Conn-Dens. | BMD | Tb.Th |
|---|---|---|---|---|---|---|
| C1 | 0.08 | 0.84 | 1.16 | 3.95 | 896.88 | 0.14 |
| C2 | 0.10 | 0.80 | 1.25 | 12.47 | 856.79 | 0.09 |
| C3 | 0.16 | 0.69 | 1.52 | 8.52 | 858.56 | 0.16 |
| C4 | 0.11 | 0.94 | 1.06 | 3.05 | 901.85 | 0.19 |
| C5 | 0.14 | 0.90 | 1.11 | 3.26 | 914.80 | 0.19 |
| C6 | 0.08 | 1.11 | 0.88 | 2.09 | 904.16 | 0.15 |
| C7 | 0.09 | 0.93 | 1.06 | 4.35 | 873.55 | 0.12 |
| C8 | 0.11 | 0.80 | 1.23 | 5.03 | 888.90 | 0.12 |
| C9 | 0.09 | 0.80 | 1.22 | 6.90 | 858.03 | 0.09 |
| C10 | 0.12 | 0.91 | 1.10 | 4.21 | 883.40 | 0.14 |
| C11 | 0.10 | 0.90 | 1.09 | 3.42 | 899.28 | 0.13 |
| C12 | 0.12 | 0.57 | 1.71 | 13.52 | 843.75 | 0.10 |
| Mean | 0.11 | 0.85 | 1.20 | 5.90 | 881.66 | 0.14 |
| STDEV | 0.02 | 0.14 | 0.22 | 3.75 | 23.01 | 0.03 |
| Mean+Sted | 0.13 | 0.99 | 1.42 | 9.65 | 904.67 | 0.17 |
| Mean-Sted | 0.09 | 0.71 | 0.98 | 2.15 | 858.66 | 0.10 |

Table S5 quantitative micro-CT scan evaluation of different PCBM phenotypes

| | | VOX-BV/TV | Tb.Sp | Tb.N | Conn-Dens | BMD | Tb.Th |
|---|---|---|---|---|---|---|---|
| Os WOT | Mean | 0.603 | 0.168 | 7.676 | 252.3 | 871.4 | 0.148 |
| | Median | 0.616 | 0.131 | 7.978 | 193.4 | 869.4 | 0.134 |
| | Sted | 0.156 | 0.089 | 2.562 | 149.0 | 33.7 | 0.049 |
| Os Tb | Mean | 0.366 | 0.349 | 4.620 | 155.3 | 844.9 | 0.129 |
| | Median | 0.341 | 0.319 | 3.832 | 102.8 | 843.0 | 0.127 |
| | Sted | 0.177 | 0.196 | 2.737 | 156.3 | 26.9 | 0.028 |
| Mix | Mean | 0.154 | 0.696 | 1.651 | 32.2 | 832.5 | 0.126 |
| | Median | 0.137 | 0.623 | 1.674 | 18.4 | 830.7 | 0.113 |
| | Sted | 0.040 | 0.194 | 0.532 | 30.7 | 24.5 | 0.030 |
| Lytic | Mean | 0.084 | 0.941 | 1.074 | 7.7 | 848.2 | 0.114 |
| | Median | 0.083 | 0.953 | 1.034 | 4.8 | 852.2 | 0.115 |
| | Sted | 0.010 | 0.106 | 0.133 | 7.5 | 23.3 | 0.022 |
| Controls | Mean | 0.108 | 0.848 | 1.199 | 5.9 | 881.7 | 0.135 |
| | Median | 0.107 | 0.867 | 1.135 | 4.3 | 886.2 | 0.134 |
| | Sted | 0.023 | 0.137 | 0.221 | 3.7 | 23.0 | 0.033 |
| P values | one-way anova | 0.0001 | 0.0711 | 0.0001 | 0.0001 | 0.0001 | 0.0001 |

Table S6

| Sample No | Lession type | Patient | VOX-BV/TV |
|---|---|---|---|
| 858 | Os Tra | 1 | 0.1611 |
| 833 | Lytic | | 0.0792 |
| 852 | Lytic | | 0.0776 |
| 857 | Mixed | 2 | 0.1239 |
| 837 | Mixed | | 0.109 |
| 843 | Mixed | 4 | 0.1384 |
| 838 | Os Tra | | 0.1318 |
| 847 | Mixed | | 0.1138 |
| 744 | Os Tra | 5 | 0.4411 |
| 747 | Os WoT | | 0.3014 |
| 821 | Os Tra | 6 | 0.1602 |
| 859 | Mixed | | 0.1098 |
| 828 | Lytic | 7 | 0.0947 |
| 799 | Lytic | | 0.0881 |
| 840 | Lytic | | 0.0744 |
| 802 | Os Tra | 10 | 0.6209 |
| 795 | Os Tra | | 0.5484 |
| 793 | Os Tra | | 0.2962 |
| 822 | Os WoT | 11 | 0.6983 |
| 823 | Os WoT | | 0.6155 |
| 830 | Os Tra | | 0.5134 |
| 869 | Os WoT | 12 | 0.7317 |
| 849 | Os Tra | | 0.5508 |
| 819 | Os WoT | | 0.4792 |
| 841 | Lytic | 15 | 0.091 |
| 798 | Lytic | | 0.0895 |
| 855 | Os Tra | 20 | 0.1953 |
| 848 | Mixed | | 0.1732 |
| 800 | Nt affeted | | 0.1129 |
| 826 | Mixed | 23 | 0.183 |
| 866 | Mixed | | 0.1081 |
| 817 | Lytic | | 0.0796 |
| 820 | Os Tra | 24 | 0.2973 |
| 816 | Mixed | | 0.1275 |
| 796 | Os Tra | 25 | 0.4163 |
| 801 | Lytic | | 0.0884 |
| 853 | Lytic | | 0.0801 |
| 794 | Os WoT | 26 | 0.7228 |
| 825 | Os WoT | | 0.6823 |
| 818 | Os WoT | | 0.3919 |
| 845 | Os WoT | 27 | 0.7913 |
| 851 | Os WoT | | 0.7599 |
| 850 | Os Tra | | 0.6627 |
| 835 | Os WoT | 28 | 0.7879 |
| 854 | Os Tra | | 0.3163 |
| 844 | Os Tra | 29 | 0.1559 |
| 839 | Mixed | | 0.1354 |
| 856 | Lytic | 40 | 0.1096 |
| 846 | Lytic | | 0.0727 |
| 815 | Os Tra | 42 | 0.6224 |
| 792 | Os WoT | | 0.6004 |
| 791 | Os Tra | 45 | 0.3799 |
| 868 | Mixed | | 0.1891 |

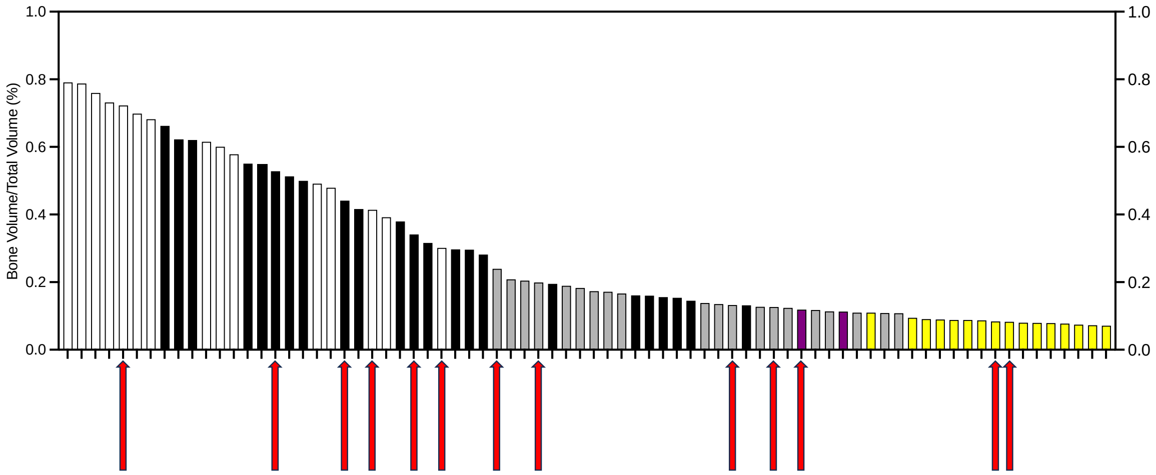


Fig S1: Waterfall plot of VB/TV of PCBM samples. Redd arrows indicates which samples were selected to perform SEM observation, and mineral composition analysis through qBSE.

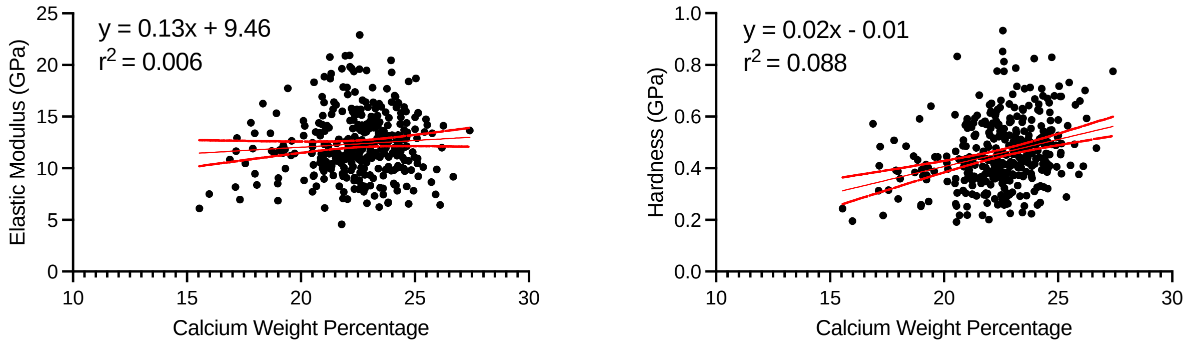


Fig S2 Linear regression between hardness and modulus with Ca content.

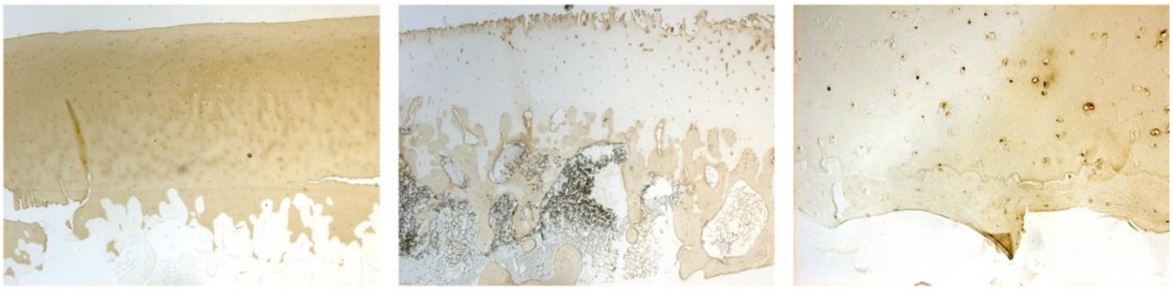

Figure S4. Immunohistochemistry in articular cartilage-bone interface. Left Collagen 2 staining, Centre Collagen 1 staining. Right aggrecan staining. Note that collagen 2 is observed in cartilage matrix but not in bone, while opposite staining is observed for Col 1, present in bone but not in catilage. Aggrecan is observed in cartilage matrix and strongly in chondrocyte lacunae.
